# Mutations that positively affect *Bandavirus* glycoprotein function on VSV vectored vaccines

**DOI:** 10.64898/2026.04.15.718665

**Authors:** Raegan J. Petch, Philip Hicks, Jonna B. Westover, Brian B. Gowen, Paul Bates

**Affiliations:** Department of Microbiology, Perelman School of Medicine, University of Pennsylvania, Philadelphia, PA, 19104; Department of Animal, Dairy, and Veterinary Sciences, Utah State University, Logan, UT, 84322

**Keywords:** bandavirus, severe fever with thrombocytopenia syndrome virus, SFTSV, Dabie bandavirus, Heartland bandavirus, vaccine, immunogenicity, neutralizing antibodies, passive transfer, rVSV, vesicular stomatitis virus, COPI

## Abstract

Severe fever with thrombocytopenia syndrome virus (SFTSV) and Heartland virus (HRTV) are emerging tick-borne bandaviruses. They have high case fatality rates (10%), and no FDA-approved vaccines exist for disease prevention. SFTSV and HRTV are therefore identified as priority pathogens. A recombinant vesicular stomatitis virus (rVSV) vaccine, which replaces the original VSV glycoprotein with the SFTSV glycoproteins, shows early promise for SFTSV as it induces strong immune responses that are protective against lethal challenge. However, rVSV-SFTSV is highly attenuated in cell culture, which may be due to incompatibility between the assembly sites of SFTSV (the Golgi and ERGIC) and that of VSV (the plasma membrane).

In this study, we identify a noncanonical COPI binding motif found in the cytoplasmic tail of SFTSV glycoproteins and demonstrate that an amino acid substitution in this motif (K1071A) inhibits binding to COPI. This mutation results in increased surface expression of SFTSV glycoproteins, improved incorporation onto VSV virions, and enhanced replication of rVSV-SFSTV *in vitro.* A mutation in a homologous site (K1074A) of HRTV has similar results, and rVSV-HRTV K1074A exhibits increased replication *in vitro* and *in vivo*. We show that vaccination with rVSV-HRTV K1074A results in improved induction of neutralizing antibody responses in immunocompetent C57BL/6 mice, and neutralizing antibodies elicited by vaccination are protective when administered to severely immunocompromised mice via passive transfer. Overall, our study identifies a mutation that improves the efficacy of the rVSV-SFTSV vaccine candidate and introduces the first vaccine candidate directly addressing HRTV infections.

**Importance:** Severe fever with thrombocytopenia syndrome virus (SFTSV) and Heartland bandavirus (HRTV) are emerging tick-borne viruses with high fatality rates. FDA-approved vaccines and antiviral drugs are unavailable but critically needed. We identify an important mutation in the SFTSV glycoprotein that disrupts a previously unreported COPI binding site. The mutation improves the efficacy of the previously described recombinant vesicular stomatitis virus vaccine candidate for SFTSV (rVSV-SFTSV). We also develop an rVSV-HRTV vaccine and show potent induction of neutralizing antibodies and protection from lethal challenge. This is the first study directly addressing the lack of vaccines specifically targeting HRTV.

## Introduction

Severe fever with thrombocytopenia syndrome virus (SFTSV, recently renamed Dabie bandavirus) and Heartland bandavirus (HRTV) are emerging tick-borne viruses in the *Bunyaviracetes* class and the *Bandavirus* genus [1–3]. SFSTV was first identified in China in 2009, and more than 30,000 cases have been recognized throughout east and southeast Asia since its discovery [1, 4–16]. Annual infections have increased in number since identification of the virus [4–6, 17], and incidences of zoonotic and human-to-human transmission have been documented with increasing frequency [18–25]. However, tick bites are still the primary mode of transmission [26]. Patients infected with SFTSV develop symptoms ranging from mild flu-like illness to severe disease including high fever, thrombocytopenia, hemorrhagic manifestations, and neurological symptoms that can progress to multiorgan failure and death [1, 5, 27]. The overall case fatality rate of SFTSV is approximately 8% [4], but regional outbreaks have had fatality rates as high as 47% [10]. Given the continued threat of SFTSV, the high mortality, and the potential for human-to-human transmission, SFTSV was designated as a priority pathogen by the World Health Organization (WHO) and the National Institute of Allergy and Infectious Diseases (NIAID) [28, 29]. Cases of SFTSV have not yet been confirmed outside of its endemic range in southeast Asia, but Heartland bandavirus (HRTV), a virus that is very closely related to SFTSV, was discovered in the United

States (US) in 2009 [2]. Since then, more than 60 cases of HRTV infection have been detected throughout the eastern US [30, 31]. However, HRTV is no longer a reportable disease in the US, and this case count is thought to be an underestimate of the true prevalence [30, 32, 33]. Seroprevalence of the virus in wildlife is high throughout the eastern US, and there are concerns that the virus is circulating more widely without detection [32–36]. Patients infected with HRTV have similar clinical signs to those with SFTSV, and the overall case fatality rate of confirmed cases is approximately 5-10% [31]. Similar to SFTSV, HRTV has been identified as a priority pathogen by NIAID [29]. Despite these designations, no vaccines or therapeutics have been approved to prevent or treat SFTSV or HRTV infections [37].

As members of the *Bunyaviracetes* class, SFTSV and HRTV have tri-segmented, single stranded RNA genomes [1, 2]. The small segment (S) is ambisense and encodes the nucleoprotein in the negative sense and a non-structural protein in the positive sense. The large (L) and medium (M) segments are negative sense and encode the RNA dependent RNA polymerase and envelope glycoproteins (GPs) respectively. The glycoprotein is translated as a polyprotein and is proteolytically cleaved into Gn and Gc [38]. Gn is predominately responsible for recognition of the receptor, resulting in binding and attachment to cells. Following attachment, Gc mediates fusion of the viral envelope with the cell membrane and subsequent entry into cells [39–43]. A definitive receptor for SFTSV has not yet been defined, but DC-SIGN, glucosylceramide, UGCG, non-muscle myosin heavy chain IIA, and CCR2 have been proposed as important entry factors [39, 40, 42, 44–46]. Receptor binding and entry is less well understood for HRTV, but C-type lectins, DC-SIGN, DC-SIGNR, LSECTin, glucosylceramide, and UGCG have been shown to be involved [44, 46, 47].

Given that the glycoproteins are responsible for attachment and entry into cells, they are the primary targets of protective immune responses, and anti-Gn neutralizing antibodies are a correlate of protection in patients infected with SFTSV [48–53]. Therefore, the majority of SFTSV vaccines in early stages of development utilize Gn and Gc as immunogens, and development of neutralizing antibodies in response to vaccination is the primary metric of success. T cell responses to SFTSV proteins in the absence of neutralizing antibodies have been shown to be protective in one vaccination study, but the role of cellular immunity in controlling natural infection is not well understood [54]. DNA, mRNA, protein subunit, whole inactivated virus, live attenuated virus, and virus-vectored vaccines have been developed for SFTSV, and the majority of these candidates provide immunity that is protective against lethal challenge in animal models [54–67]. However, almost all of the candidates require at least 2 doses for complete immunity, which is a hindrance to effective vaccination in rural areas where tick exposure is most common and vaccination is most urgently needed [68, 69]. The recombinant vesicular stomatitis virus vaccine (rVSV-SFTSV) has unique advantages because it only requires one dose for complete protection and maintains a positive safety profile [63, 64]. rVSV-SFTSV has also been shown to have efficacy in cross-protection against lethal HRTV challenge, which is of high importance given that no studies directly evaluating HRTV vaccines have been published [64].

Vesicular stomatitis virus (VSV) is a rhabdovirus that has been developed as a vaccine vector due to its ability to incorporate foreign glycoproteins onto virions and induce strong, protective antibody and T cell responses against the foreign antigen [70]. An FDA-approved recombinant VSV vaccine for Zaire Ebola virus (rVSV-ZEBOV) has been used with remarkable success. A single dose of rVSV-ZEBOV results in rapid development of long-lived protective antibody and T cell responses [71–74]. Importantly, rVSV-ZEBOV was also shown to be protective if given shortly after exposure in animal studies and some cases of incidental human exposure, which is an attribute that few other vaccines have achieved [75–81]. The safety of rVSV-ZEBOV is well documented, and it has now been approved for use in children [82]. The incredible success of the rVSV-ZEBOV vaccine sets a precedent for the use of this platform for emerging diseases, including SFTSV and HRTV.

rVSV-SFTSV is a promising vaccine candidate, but further studies are required to optimize the virus and enhance our understanding of its therapeutic value. rVSV-SFTSV is attenuated in cell culture, and data from our lab have shown that titers of rVSV-SFTSV are approximately 90-fold lower than those of the parent VSV at 24 hours post infection [64]. Recovery of rVSV-SFTSV through plasmid-based reverse genetics systems is also challenging and is much less efficient than that of unaltered VSV or other rVSV encoding foreign glycoproteins such as ZEBOV. rVSV-SFTSV attenuation may be due to inefficient incorporation of SFTSV glycoproteins onto VSV particles due to incompatibility in viral assembly sites. VSV assembles at the cell membrane, but SFTSV glycoproteins are targeted to the endoplasmic reticulum (ER)-Golgi intermediate complex (ERGIC) and Golgi for assembly and egress [83].

Like many other bunyaviruses, SFTSV Gn distributes to the Golgi when expressed independently of Gc while Gc exhibits an ER distribution pattern [84–91]. Signals in Gn are considered to dictate the localization of the Gn/Gc heterodimer since expression of Gn is sufficient to recruit Gc to the Golgi from the ER. Protein sorting signals responsible for these localization patterns are loosely recognized based on studies of other bunyaviruses, but mechanisms specific to SFTSV are not well understood. Golgi targeting signals have been mapped to the Gn cytosolic tail, transmembrane domain, or both the cytosolic tail and transmembrane domain of genetically related bunyaviruses, but generalizations to SFTSV are undermined by poor sequence conservation within these regions [88, 91–93]. ER retention of bunyaviral Gc has been attributed to the C-terminus of the cytosolic tail [93–95], which is enriched in basic residues that fit or approximate well-established KxKxx or KKxx (where K is lysine and x is any amino acid) motifs known to bind coatomer complex I (COPI) [96–98]. COPI participates in retrograde trafficking between the Golgi cisternae and from the Golgi to the ER [99]. Previous studies have implicated or confirmed the role of COPI in the intracellular retention of some coronaviral spike glycoproteins [100–106], but direct investigation of the role of COPI in bunyavirus replication is generally lacking.

We hypothesize that the attenuation of rVSV-SFTSV may impair replication and immunogenicity *in vivo.* Data from our lab and others have identified four mutations (C617R, M749T, E982K, and K1071E) that arise spontaneously in SFTSV Gc upon passage of rVSV-SFTSV and may increase the replication of the rVSV-SFTSV virus in cell culture [64, 107]. However, the mechanisms by which these mutations increase rVSV-SFTSV replication are not well understood, and synergism between these mutations has not yet been evaluated. While spontaneous acquisition of mutations may be a positive outcome for *in vitro* replication, it may present a challenge for clinical translation of the vaccine candidate due to concerns about instability of the viral genome. Moreover, spontaneous mutations may inadvertently disrupt epitopes important for immune control of circulating viruses. It is therefore favorable to use reverse genetics systems to genetically alter the SFTSV GP on rVSV to introduce stable mutations with known biological outcomes and predictable immunogenicity.

In this study, we evaluate the mutations C617R, M749T, E982K, and K1071A in the context of the SFTSV glycoprotein. We found that the combination of K1071A+M749T, K1071A+E982K, or K1071A+C617R+M749T results in increased titers of VSVΔG(SFTSV) pseudotypes. Evaluation of the mechanisms responsible shows increased stability, fusogenicity, and surface expression of the mutant SFTSV glycoproteins. The increased surface expression is striking and is caused solely by the K1071A substitution. As seen with other bunyaviruses [93, 94], we find that this mutation is sufficient to relocalize SFTSV glycoproteins from intracellular compartments to the cell surface. We also use immunoprecipitation assays to demonstrate that WT SFTSV Gc binds to β’-COP, and that the K1071A mutation decreases the strength of this interaction. To our knowledge, this is the first report providing direct evidence that COPI binding is important for bunyavirus protein localization. Furthermore, we establish that the K1071A mutation increases the efficiency of replication of rVSV-SFTSV and improves the incorporation of the SFTSV glycoprotein onto VSV virions. Mutations made in homologous sites in rVSV-HRTV result in a similar phenotype, where the same mutation in the cognate position of HRTV Gc (K1074A) also improves viral replication and surface glycoprotein expression in infected cells. Finally, we evaluate the immunogenicity of the rVSV-HRTV K1074A vaccine and demonstrate that it improves production of neutralizing antibody responses in C57BL/6 mice and exhibits increased *in vivo* replication in immunodeficient AG129 mice. We also show that neutralizing antibodies developed in response to rVSV-HRTV K1074A are sufficient to protect naïve AG129 mice from lethal challenge when administered via passive transfer.

## Materials and Methods

### Plasmids

Cloning was performed using PCR products amplified by Q5 Hot Start High-Fidelity DNA Polymerase (New England Biolabs, M0494). PCR products were purified from agarose gels using Monarch DNA Gel Extraction Kit (New England Biolabs, T1020) using manufacturer’s protocol and ligated using NEBuilder HiFi DNA Assembly Master Mix (New England Biolabs, E2621). All plasmids were grown in NEB 5-alpha Competent *E. coli* (New England Biolabs, C2987) and purified using ZymoPURE Plasmid Miniprep Kit (Zymo research, D4210) or GenElute HP Plasmid Maxiprep Kit (Sigma-Aldrich, NA0310-1KT). All plasmids were submitted for Nanopore sequencing prior to use (Eurofins Genomics).

*Cloning of plasmids encoding full-length bunyavirus glycoproteins:* Full-length DNA sequences encoding the codon-optimized coding sequences (gBlocks, Integrated DNA Technologies) of the M segments from SFTSV and HRTV were amplified by PCR and cloned into the mammalian expression plasmid pCG1 to produce pCG1-SFTSV-Gn/Gc and pCG1-HRTV Gn/Gc. Site directed mutagenesis primers were designed to amplify the entirety of pCG1 and contain mutations encoding C617R, M749T, E982K, or K1071A substitutions for SFTSV or C621S, V753T, or K1074A substitutions for HRTV. In our original study, we determined that a mutation encoding K1071E substitution occurred in the SFTSV glycoprotein upon passage of rVSV-SFTSV, but we decided to evaluate a K1071A substitution instead based on other reports of this substitution in similar positions of Rift Valley fever virus (RVFV), Uukuniemi virus (UUKV), and SARS-CoV-2, among others [93, 94, 100]. Similarly, we chose to evaluate a C621S substitution in HRTV as opposed to the C621R substitution based on reports that a C621S substitution arises upon passage of HRTV in mice [108]. The single PCR products for each plasmid were ligated by NEBuilder HiFi DNA assembly. Following the production of plasmids with each mutation alone, we used the same mutagenesis primers to produce plasmids with multiple mutations. In total, we produced pCG1-SFSTV Gn/Gc WT; C617R; M749T; E982K; K1071A; C617R+M749T; K1071A+C617R; K1071A+M749T; K1071A+E982K; and K1071A+C617R+M749T. We also produced pCG1-HRTV Gn/Gc WT; K1074A; and C621S+V753T+K1074A.

*Cloning of pVSV-SFTSV launch plasmids:* The pVSV eGFP RABV-G plasmid (Addgene, #31833) was used as the template for creating vectors expressing the VSV antigenome with SFTSV or HRTV glycoproteins in place of the VSV-G glycoprotein. Primers were designed to amplify the pVSV vector backbone with overlaps matching the 5’ and 3’ ends of SFTSV or HRTV glycoproteins. The SFTSV or HRTV glycoprotein (WT or with the indicated substitutions) was amplified from the pCG1 plasmids described above using primers designed to overlap with the pVSV vector backbone. NEBuilder HiFi DNA Assembly was performed to ligate the glycoprotein inserts to the pVSV vector.

*Cloning of VSV helper plasmids for VSV launch:* The pCG1 expression vector was modified to remove two T7 promoters, which were hypothesized to compete for T7-driven polymerase expression of the pVSV vector plasmid, to produce the expression plasmid pCG2. Full length, codon-optimized sequences of VSV-N, -P, and - L were synthesized (gBlocks, Integrated DNA Technologies) based on previous reports [109] and cloned into pCG2 expression vector downstream of the rabbit beta-globin intron. T7opt in the pCAGGS vector was a gift from Benhur Lee (Addgene, #65974) [110].

*Cloning of fluorescent SFTSV glycoprotein chimeras:* Sequences encoding mCherry (amino acids 2-236) and mNeonGreen (amino acids 2-236) were amplified by PCR. PCR using pCG1-SFTSV-Gn/Gc WT or K1071A template DNA was performed to delete sequences encoding the ectodomain of Gc (amino acids 567-1029). NEBuilder HiFi DNA Assembly was performed using reactions containing two products to generate the cloning intermediate plasmids pCG1-SFTSV-Gn/Gc^ΔEcto/mCherry^ and pCG1-SFTSV-Gn/Gc^ΔEcto/mNeonGreen^. To generate pCG1-SFTSV-Gc^ΔEcto/mCherry^ and pCG1-SFTSV-Gc^ΔEcto/mNeonGreen^, PCR was performed on pCG1-SFTSV-Gn/Gc^ΔEcto/mNeonGreen^ and pCG1-SFTSV-Gn/Gc^ΔEcto/mCherry^ to generate a product in which the sequence encoding the entirety of Gn (amino acids 1-535) was deleted. These products were circularized using NEBuilder HiFi DNA Assembly. These plasmids are hereby referred to as mNeonGreen-RxKxx, mNeonGreen-RxAxx, mCherry-RxKxx, and mCherry-RxAxx to reflect plasmids in which the ectodomain of Gc was replaced by a fluorescent reporter appended to either the WT transmembrane domain and cytoplasmic tail (-RxKxx) or the transmembrane domain and cytoplasmic tail with a lysine to alanine mutation in the −3 position (-RxAxx).

### Cells

ATCC verified and mycoplasma free HEK293T (293T), Vero E6, A549, and HeLa cells were maintained in DMEM (Corning, MT10-013-CV) with 10% cosmic calf serum (CCS) (HyClone, #SH30087.03). Cells were passaged every 2-3 days using Trypsin-EDTA (0.05%) (Invitrogen, 25300054). Where indicated, cells were seeded in plates coated with 0.025 mg/mL rat tail collagen (Corning, 354236) in 0.02M acetic acid.

To launch rVSV-SFTSV, we developed HEK293T cells that stably expressed CCR2, which is known to be an important entry factor for SFTSV [42]. A plasmid encoding human CCR2 transcript variant B was purchased from GenScript (OHu23522) and was transfected into HEK293T cells using Lipofectamine 3000 according to manufacturer’s guidelines. One day after transfection, the cells were passaged into media containing 500 μg/mL Geneticin (Gibco, 10131-035). Surviving colonies were passaged and expanded in media with Geneticin.

### Production of VSVΔG(SFTSV) Mutant Pseudotypes

VSVΔG(SFTSV) pseudotypes were produced as previously described [111]. Briefly, 293T cells were seeded in collagen-coated 10-cm plates in the afternoon to achieve a density of 70% confluency the following morning. Cells were transfected with 25 μg of purified DNA from plasmids encoding the SFSTV mutant glycoproteins using calcium phosphate according to the manufacturer’s protocol (Takara Bio, 631312). Approximately 24 hours post transfection (hpt), transfected cells were infected with VSVΔG-RFP virus pseudotyped with VSV-G at a multiplicity of infection of ∼1-3 at 37⁰C. Approximately 24 hours post infection (hpi), supernatant was harvested from infected cells and clarified twice by centrifugation at 1250×g for 10 min at 4°C. Purified supernatant was aliquoted and stored at −80°C.

To titer the pseudotypes, Vero E6 cells were seeded in the afternoon in 100 μL at a density of 2.5×10^4^ cells/well into a collagen-coated 96-well plate. The next morning, VSVΔG-RFP(SFTSV) mutant pseudotypes were serially diluted 2-fold in DMEM+10% CCS. Dilution medium also contained 600 ng/mL anti-VSV-G monoclonal antibody 1E9F9, which was used to neutralize any carryover pseudovirus containing VSV-G. The virus/antibody mixtures were incubated at 37°C for 1 hour before being transferred onto Vero E6 cells. 30 hours post-infection, cells were gently washed with PBS and fixed with 4% paraformaldehyde. Transduced cells were detected by RFP fluorescence using an S6 FluoroSpot Analyzer (CTL, Shaker Heights, OH). Individual spots were counted using Basic Count settings with spot sizes. Average foci counts from uninfected wells were subtracted from all wells on the same plate to correct for background. Infectious titers (focus forming units, FFU/mL) were calculated per sample by averaging values collected from all wells within the linear range on the dilution curve.

### SFSTV Mutant Glycoprotein Surface Expression Flow Cytometry

A549 cells were seeded into 6-well plates in the afternoon to achieve a density of 80% confluency the following morning. Cells were transfected with 2 μg of purified SFTSV mutant glycoprotein DNA using Lipofectamine 3000 according to the manufacturer’s protocol (Invitrogen, L3000-015). Cells were incubated in Opti-MEM (Gibco, 31985070) following transfection and supplemented with pre-warmed DMEM+10% CCS 3 hpt. Transfected cells were lifted off the plate 24 hpt using PBS with 10 mM EDTA. Cells were pelleted at 400×g for 5 minutes at 4°C and resuspended in FACS buffer (PBS with 2% BSA and 0.1% sodium azide) containing LIVE/DEAD Fixable Violet Dead Cell Stain (Invitrogen, L34964) at 1:40 dilution and serum from mice immunized with an mRNA LNP encoding the unaltered, full-length SFTSV Gn/Gc [112] at a dilution of 1:1000. Cells were incubated in this LIVE/DEAD and primary antibody mixture in the dark for 1 hour at room temperature to label proteins on the cell surface. Following incubation, cells were pelleted and washed 3 times with FACS buffer, then fixed with 4% paraformaldehyde for 20 minutes. Cells were briefly washed twice with FACS buffer, and then they were permeabilized with FACS buffer containing 0.1% Triton X-100 for 30 minutes. To label intracellular proteins, cells were incubated for an additional hour with a mixture of primary rabbit polyclonal antibodies against SFTSV Gn (ProSci, #6647) and SFTSV Gc (ProSci, #6653), each at a 1:2000 dilution. Cells were again washed three times with FACS buffer, and then they were stained with 1:1000 dilutions of donkey anti-mouse IgG Alexa Fluor 488 (Invitrogen, A21202) and donkey anti-rabbit IgG Alexa Fluor 594 (Invitrogen, A21207) for 1 hour at room temperature. Cells were washed three final times with FACS buffer and analyzed on a BD LSRII flow cytometer with high-throughput system using FACSDIVA software (BD Biosciences). Data analysis was performed using FloJo software (FloJo LLC). Gating was performed to select live single cells based on low violet fluorescent reactive dye staining. Stained mock-transfected cells were used to gate for both surface-positive and permeabilized-positive signal. The MFI of surface-positive (AF488) and permeabilized-positive (AF594) antigen was calculated, and the surface-positive MFI was compared to the MFI of total antigen (surface-positive + permeabilized-positive) to quantify the percentage of SFSTV antigens that were surface expressed.

### SFTSV Mutant Glycoprotein Stability

293T cells were seeded into collagen-coated 6-well plates in the afternoon to achieve a density of 80% confluency the following morning. Cells were transfected in triplicate with 2 μg of purified SFTSV mutant glycoprotein DNA using Lipofectamine 3000 according to the manufacturer’s protocol (Invitrogen, L3000-015). Cells were incubated in Opti-MEM (Gibco, 31985070) following transfection and supplemented with pre-warmed DMEM+10% CCS 3 hpt. Forty-eight hpt, two wells of cells transfected with each glycoprotein were treated with cycloheximide (Sigma-Aldrich, C7698-1G) at a final concentration of 50 mg/mL according to established protocols [113]. The third well of each condition was treated with DMSO as a vehicle control.

Six hours post treatment, one well of cycloheximide treated cells and the DMSO treated cells were harvested for each SFTSV mutant. The cells were washed gently with ice cold PBS and removed from the plate. They were pelleted at 400×g for 5 minutes at 4°C and resuspended in 200 μL Triton X-100 cell lysis buffer with cOmplete Protease Inhibitor Cocktail (Millipore Sigma, 11836170001). Cells were incubated at 4°C for lysis, and then debris was pelleted by centrifuging at 15,000 rpm for 10 minutes at 4°C. The supernatant was transferred to a clean tube and stored at −20°C prior to analysis via Western Blot. The above protocol was repeated for the remaining well at 10 hours post treatment.

Protein samples were quantified via Nanodrop and normalized to the same concentration prior to Western Blot. Laemmli sample buffer was added, and the proteins were denatured at 95°C for 5 min. Samples were run on a 4-15% Criterion Tris-HCl Protein Gel (Bio-Rad Laboratories, 3450028). Protein was transferred to an Amersham Hybond P 0.45 PVDF membrane (Cytiva, 10600029) and blocked in 5% milk in TBS buffer with 0.1% Tween-20 (TBS-T) for one hour. Proteins were probed with rabbit polyclonal anti-SFTSV Gc antibody at a 1:5000 dilution (Pro Sci, #6651) and a mouse monoclonal anti-hGAPDH antibody at 1:500 dilution (Developmental Studies Hybridoma Bank, DSHB-GAPDH-2G7) overnight at 4°C. The membranes were washed three times with TBS-T and then incubated with HRP-conjugated secondary antibodies (Invitrogen 31430 – 1:100,000 and A27036 – 1: 30,000) for 20 minutes at room temperature. The membranes were again washed three times with TBS-T and then developed using SuperSignal West Femto Maximum Sensitivity Substrate (Thermo Scientific, 34095) and read on a GE Healthcare Amersham 600 imager. Band intensity was analyzed via FIJI (ImageJ).

### SFSTV Mutant Glycoprotein Fusogenicity

HeLa cells were seeded into collagen-coated 6-well plates in the afternoon to achieve a density of 80% confluency the following morning. Cells were transfected with 2 μg of purified SFTSV mutant glycoprotein DNA using Lipofectamine 3000 according to the manufacturer’s protocol (Invitrogen, L3000-015). Cells were incubated in Opti-MEM (Gibco, 31985070) following transfection and supplemented with pre-warmed DMEM+10% CCS 3 hpt. Twenty-four hpt, cells were removed from the plates using Trypsin-EDTA (0.05%) (Invitrogen, 25300054) and seeded in new collagen-coated 6-well plates to achieve a monolayer the following day. The next day (48 hpt), media was removed from the cells, and they were washed twice with PBS to remove all media. They were treated with PBS buffered with 50 mM MES monohydrate (Thermo Scientific, A16104.22) adjusted to pH 5.5 for 5 minutes. Following the low pH incubation, the acidic buffer was removed, the cells were washed twice more with PBS, and then 2 mL of pre-warmed DMEM+10% CCS was added to the cells. They were incubated for 4 hours, and then brightfield images of five high-powered fields per condition were taken. Syncytia per high-powered field and nuclei per syncytia were counted for each condition.

### Fluorescence Microscopy

A549 cells were seeded onto collagen-coated 96-well black plates with #1.5 high performance cover glass (0.17 ± 0.005mM) bottoms in the afternoon. The following morning, cells at a confluency of ∼90% were transfected with purified plasmid DNAs using Lipofectamine 3000 (Invitrogen, L3000-015) according to the manufacturer’s protocol. Media was replaced on cells 6 hours post-transfection (hpt). Cells were gently washed with PBS 24 hpt and were subsequently fixed with 4% paraformaldehyde. In indicated experiments, cells were treated with 33 μM nocodazole (Thermo Scientific, AC358240100) for 3 hours until cells were washed and fixed 24 hpt.

To stain cells for immunofluorescence microscopy, fixed cells were first blocked for 1 hour in blocking buffer (PBS with 0.1% sodium azide and 10% heat-inactivated donkey serum (Sigma, D9663-10mL)) at room temperature. In assays specifically analyzing surface antigen, unpermeabilized cells were covered in 50 μL SFTSV-vaccinated mouse serum diluted 1:5000 in staining buffer (PBS with 0.1% sodium azide and 5% donkey serum) and incubated for 1 hour at room temperature. Cells were washed three times with staining buffer before being fixed again with 4% paraformaldehyde and permeabilized with blocking buffer containing 0.1% Triton X-100 for 30 min at room temperature. Intracellular antigen was labelled by staining with anti-SFTSV Gc rabbit polyclonal antibody (Pro Sci, #6653) diluted 1:1000 in staining buffer for 1 hour at room temperature. In indicated experiments, β’-COP was labeled with a polyclonal rabbit antibody at a dilution of 1:500 (Invitrogen, PA5-96557). Cells were washed three times and then stained with secondary antibodies (Thermo Scientific, A-21125 and A21206) at 1:1000 dilution for 1 hour at room temperature. All cells were stained using 10 μM Hoechst 33258 (Enzo Life Sciences, ENZ-52402) diluted in PBS for 30 min. Images were captured on a Nikon Eclipse Ti2-E inverted microscope (Nikon, Minato City, Tokyo, Japan) using an ORCA-flash4.0 V3 digital CMOS camera (Hamamatsu Photonics K.K., Hamamatsu City, Japan, #C13440). Imaging at 100x magnification was performed with immersion oil type F (Nikon, MXA22168). Laser intensities and exposure times were held constant across a single experiment to permit downstream quantitative analysis. Images were deconvolved using NIS-Elements software (Nikon, Minato City, Tokyo, Japan) with automatic settings.

Images were analyzed and prepared for publication using FIJI (ImageJ) software [114]. Brightness/contrast were adjusted and held constant across a single experiment. Colocalization was quantified by fractional overlap on representative equatorial slices of z-stack images [115]. Briefly, a region of interest was drawn around each individual cell. To perform background subtraction, pixel intensity was measured in the nuclear region of each cell, and median pixel intensity was subtracted globally from each ROI using the Subtract command. This process was performed iteratively until the median pixel intensity for the nuclear region was 0. Channels were split and an automatic local threshold was applied to each with a Bernsen radius of 5. Channel images were then converted to binary masks. Binary images of two channels were multiplied together using the Image Calculator to create a binary image of overlapping pixels. Signal area of all relevant binary images was measured using the Analyze Particles command and size inclusion criteria of 5 to infinity pixels. Fractional overlap coefficients were calculated by dividing total overlapping pixel area by the pixel area of the channel of interest.

### mCherry Flow Cytometry

The surface expression of mCherry-RxKxx, mCherry-RxAxx, or cytosolic mCherry was evaluated using flow cytometry, similar to the methods described above. Briefly, A549 cells were seeded in 6-well plates in the afternoon to achieve a density of 80% confluence the following morning. Cells were transfected with 1 μg of purified mCherry-RxKxx, mCherry-RxAxx, or cytosolic mCherry plasmid DNA using Lipofectamine 3000 as described above. Transfected cells were harvested 24 hpt using PBS with 10 mM EDTA. Cells were pelleted at 400×g for 5 minutes at 4°C and resuspended in FACS buffer (PBS with 2% BSA and 0.1% sodium azide) containing LIVE/DEAD Fixable Violet Dead Cell Stain (Invitrogen, L34964) at 1:20 dilution for 30 minutes. Following incubation, cells were pelleted and washed 3 times with FACS buffer, then fixed with 4% paraformaldehyde for 20 minutes. Cells were briefly washed twice with FACS buffer and then were resuspended in FACS buffer containing a mouse monoclonal antibody against mCherry (BioLegend, 677702) and donkey anti-mouse IgG Alexa Fluor 488 (Invitrogen, A21202) for 2 hours. Cells were washed three final times with FACS buffer and analyzed on a BD LSRII flow cytometer with high-throughput system using FACSDIVA software (BD Biosciences). Data analysis was performed using FloJo software (FloJo LLC). Gating was performed to select live single cells based on low violet fluorescent reactive dye staining. The baseline autofluorescence of mock-transfected cells was used to subtract background fluorescence from all conditions. The MFI of surface-positive (AF488) and intracellular (mCherry) antigen was calculated, and the surface-positive MFI was compared to the MFI of total antigen (surface-positive + intracellular-positive) to quantify the percentage of SFSTV antigens that were surface expressed.

### Β’-COP Immunohprecipitation

293T cells were seeded in collagen-coated 10-cm plates in the afternoon to achieve a density of 80% confluence the following morning. Cells were transfected with 5 μg of purified SFTSV mutant glycoprotein DNA and 5 μg of purified β’-COP DNA with a C-terminal FLAG tag (GenScript, OHu04585D) using Lipofectamine 3000 according to the manufacturer’s protocol (Invitrogen, L3000-015). Cells were incubated in Opti-MEM (Gibco, 31985070) following transfection and supplemented with pre-warmed DMEM+10% CCS 3 hpt. Forty-eight hpt, the cells were washed gently with ice cold PBS and removed from the plate. They were pelleted at 400×g for 5 minutes at 4°C and resuspended in Triton X-100 cell lysis buffer with cOmplete Protease Inhibitor Cocktail (Millipore Sigma, 11697498001). Cells were incubated at 4°C for lysis, and then debris was pelleted by centrifuging at 15,000 rpm for 10 minutes at 4°C. The supernatant was transferred to a clean tube and stored at −20°C prior to analysis via Western Blot and immunoprecipitation.

For immunoprecipitation, 1 mL of lysate containing the SFTSV mutant glycoprotein and the FLAG-tagged β’-COP plasmid was transferred to a separate tube containing 100 μL of Pierce Anti-DYKDDDDK Magnetic Agarose (Thermo Scientific, A36797). The tubes containing the lysate and the magnetic beads were incubated on an inverter overnight at 4°C. The rest of the lysate was used for the “input” Western blot to ensure that equal amounts of protein were expressed for each condition. Following overnight incubation, beads were isolated using a magnetic rack and washed three times with Triton X-100 lysis buffer containing cOmplete Protease Inhibitor Cocktail (Millipore Sigma, 11697498001). 70 μl of Laemmli sample buffer was added to the beads for each condition, and proteins were denatured at 95°C for 5 minutes to elute the protein from the beads.

Immunoprecipitation samples were run on a 4-15% Criterion Tris-HCl Protein Gel (Bio-Rad Laboratories, 3450028). Protein was transferred to an Amersham Hybond P 0.45 PVDF membrane (Cytiva, 10600029) and blocked in 5% milk in TBS buffer with 0.1% Tween-20 (TBS-T) for one hour. Proteins were probed with rabbit polyclonal anti-SFTSV Gc antibody at a 1:5000 dilution (Pro Sci, #6651) overnight at 4°C. The membranes were washed three times with TBS-T and then incubated with a highly cross-adsorbed HRP-conjugated anti-rabbit secondary antibody (Invitrogen A16110) at a dilution of 1:20,000 for 20 minutes at room temperature. The membranes were again washed three times with TBS-T and then developed using SuperSignal West Femto Maximum Sensitivity Substrate (ThermoScientific, 34095) and read on a GE Healthcare Amersham 600 imager. Input samples were run in the same way but were stripped with Restore PLUS Western Blot Stripping Buffer (Thermo Scientific, 46430) after the SFTSV probe. They were reprobed with a rabbit β’-COP polyclonal antibody (Invitrogen, PA5-96557) at a 1:6000 dilution and then washed and incubated with the highly cross-adsorbed HRP-conjugated antibody as described above.

### rVSV-SFTSV and rVSV-HRTV Launch

Recombinant VSV-SFTSV and VSV-HRTV viruses with an additional open reading frame encoding eGFP in the first position were launched as described previously [109]. Briefly, 293T cells that were designed to stably express CCR2 were seeded in collagen-coated 6-well plates in the afternoon to reach a density of 60% confluence the following morning. Cells were transfected with purified DNA from plasmids encoding the VSV genome, VSV-N, VSV-P, VSV-L, and T7 polymerase at a 1:3:1:1:2 molar ratio with a total of 2.7 μg DNA per well. Transfections were done with Lipofectamine 3000 according to the manufacturer’s protocol (Invitrogen, L3000-015). Cells were incubated in Opti-MEM (Gibco, 31985070) following transfection and supplemented with pre-warmed DMEM with 10% CCS 3 hours post-transfection. Cells were watched closely over a period of 5-7 days, and media was replaced with pre-warmed DMEM+10% CCS whenever the media appeared spent. Cells were observed for eGFP expression and cytopathic effect (CPE) including cell rounding and death. Supernatant was harvested from cells once most cells were infected and clarified via centrifugation at 1250×g for 10 minutes at 4C, then clarified supernatant was passaged onto Vero E6 cells to produce passage 1 of each virus.

Large stocks of each rVSV-SFTSV and rVSV-HRTV virus were grown in Vero E6 cells by infecting a confluent 15-cm plate at an MOI of 0.0001. Virus-containing supernatants were collected 48-72 hpi and were clarified by centrifuging twice at 1250×g for 5 min at 4°C. Supernatants were then passed through a syringe filter with 0.22 μm pore size (Corning, 431219) to remove cellular debris. Virus was then frozen at −80°C until used for ultracentrifugation. Virus was concentrated by ultracentrifugation of virus-containing media through a sucrose cushion at 26,000 rpm for 2 h at 4 °C using SW-32 tubes in a Beckman Coulter Optima XPN-80 ultracentrifuge (Beckman Coulter). After removal of the sucrose and media, pelleted virus was placed on ice with 1 mL sterile TBS per 15-cm plate overnight. The next day, virus pellets were gently resuspended, aliquoted, and frozen at −80 °C. Viral titer was determined by plaque assays on Vero E6 cells with a 1.25% Avicel RC-591 NF (DuPont, #RC591-NFBA500) overlay and then stained with 1% crystal violet. To assess the genetic stability of the glycoproteins, all rVSV-SFTSV and rVSV-HRTV viruses were passaged on Vero E6 cells for a minimum of 5 passages. Viral RNA was collected and extracted, and cDNA from each sample was amplified via PCR and sequenced via Nanopore sequencing technology (Eurofins Genomics).

Replication kinetics were evaluated via a standard replication time course experiment. Briefly, Vero E6 cells were seeded in 10-cm plates in the afternoon to achieve a monolayer the following morning. The next day, the cells were infected at an MOI of 0.001. Supernatant samples were collected at 12-, 24-, 48-, and 72-hours post infection, and the media removed was replaced with an equivalent volume of pre-warmed DMEM with 10% CCS. Viral titers of each virus at each timepoint were quantified via plaque assays on Vero E6 cells. Plaques produced by each virus were measured 96 hpi with a standard ruler and rounded to the nearest quarter of a mm.

### Glycoprotein Surface Expression in rVSV-SFTSV and rVSV-HRTV Infected Cells

Similar to the methods described above, the surface expression of glycoproteins in rVSV-SFTSV and rVSV-HRTV infected cells was evaluated by flow cytometry. A549 cells were seeded in collagen-coated plates in the afternoon to achieve a monolayer the following morning. Cells were infected with the rVSV-SFTSV and rVSV-HRTV mutants at an MOI of 2. rVSV-SFTSV M749T and rVSV-HRTV WT (p2) could not be concentrated to a titer that would permit infection at the same MOI, and the replication kinetics of these viruses prohibited analysis at the same time point as the other viruses. For this reason, these samples were excluded from these analyses. Twenty-four hours after infection, samples were harvested using PBS with 10 mM EDTA and prepared for flow cytometry. As described above, cells were pelleted at 400×g for 5 minutes at 4°C and resuspended in FACS buffer containing LIVE/DEAD Fixable Violet Dead Cell Stain (Invitrogen, L34964) at 1:40 dilution and serum from mice vaccinated against SFSTV glycoproteins or HRTV glycoproteins. Cells were incubated in this LIVE/DEAD and primary antibody mixture in the dark for 1 hour at room temperature to label proteins on the cell surface. Following incubation, cells were pelleted and washed 3 times with FACS buffer, then fixed with 4% paraformaldehyde for 20 minutes. Cells were briefly washed twice with FACS buffer, and then they were stained with 1:1000 dilution of donkey anti-mouse IgG Alexa Fluor 594 (Invitrogen, A21203) for one hour at room temperature. Cells were washed three final times with FACS buffer and analyzed on a BD LSRII flow cytometer with high-throughput system using FACSDIVA software (BD Biosciences). Data analysis and flow cytometry plot construction were performed using FloJo software (FloJo LLC). Gating was performed to select live single cells based on low violet fluorescent reactive dye staining and infected cells via positive GFP staining. Stained mock-infected cells were used to gate for both infected (GFP) and surface antigen-positive (Alexa Fluor 594) signal. The MFI of surface-positive antigen in all infected cells was calculated to quantify the total surface expression of SFTSV or HRTV glycoproteins in infected cells.

### Quantification of Glycoprotein Incorporation into Virions

Western Blot analysis was used to quantify SFTSV and HRTV glycoprotein incorporation into purified VSV virions. Triton X-100 lysis buffer with cOmplete Protease Inhibitor Cocktail (Millipore Sigma, 11697498001) was added directly to purified virions. Laemmli sample buffer was added, and the proteins were denatured at 95°C for 5 min. Samples were run on a 4-15% Criterion Tris-HCl Protein Gel (Bio-Rad Laboratories, 3450028). Protein was transferred to an Amersham Hybond P 0.45 PVDF membrane (Cytiva, 10600029) and blocked in 5% milk in TBS buffer with 0.1% Tween-20 (TBS-T) for one hour. Proteins were probed with rabbit polyclonal anti-SFTSV Gc antibody at a 1:5000 dilution (Pro Sci, #6651), rabbit polyclonal anti-SFTSV Gn antibody at a 1:3000 dilution (Pro Sci, #6647) and a mouse monoclonal anti-VSV-M antibody at 1:5000 dilution (Kerafast, 23H12). The SFTSV Gn and Gc antibodies were also used to assess incorporation of HRTV glycoproteins into VSV virions because they demonstrated cross-reactivity against HRTV glycoproteins. Membranes were stripped with Restore PLUS Western Blot Stripping Buffer (Thermo Scientific, 46430) between each antibody. Proteins were detected with HRP-conjugated secondary antibodies (Invitrogen 31430 – 1:100,000 and A27036 – 1: 30,000) for 20 minutes at room temperature, and the membranes were developed using SuperSignal West Femto Maximum Sensitivity Substrate (Thermo Scientific, 34095) and read on a GE Healthcare Amersham 600 imager. Band intensity was analyzed via FIJI (ImageJ).

### rVSV-HRTV Vaccination Studies

To evaluate neutralizing antibody responses to rVSV-HRTV in immunocompetent mice, we vaccinated five male and five female eight-week-old C57BL/6 mice (The Jackson Laboratory, #000664). The mice were anesthetized with isoflurane and vaccinated intraperitoneally (IP) with 10^6^ PFU of rVSV-HRTV WT (p3), K1074A, or C621S+V753T+K1074A at day 0 and again at day 28. The viruses were diluted in sterile PBS immediately prior to injection, and the total injection volume was 150 μL. We collected blood from mice anesthetized with isoflurane via the submandibular route, and the blood was processed immediately following collection. Briefly, serum was isolated via centrifugation at 8,000 RPM for 30 minutes at 4°C in an Eppendorf 5424R centrifuge (Eppendorf, Enfield, CT). EDTA was added to each sample for a final concentration of approximately 5mM. Serum was heat inactivated by incubating at 53°C for 30 minutes, then centrifuged at 15,000 RPM for 20 minutes. Serum samples were stored at −20 °C.

To evaluate protection from lethal challenge, eight-week-old AG129 mice (deficient in interferon α/β and γ receptors) were obtained from breeding colonies at Utah State University. Six male and six female mice per group were IP vaccinated with 10^2^ PFU of each rVSV-HRTV vaccine in a 100 μL inoculum. Again, the vaccines were diluted in sterile PBS immediately prior to injection. An additional control group of four male and four female mice was injected with 100 μL PBS. Mice were weighed and closely evaluated for signs of illness each day following vaccination, and approximately 50 μL of blood was collected from a subset of mice in each group two days post vaccination to quantify rVSV replication via qPCR. Twenty-one days post vaccination, 200 μL of blood was collected for neutralizing antibody analysis, and serum was isolated as indicated above. Twenty-three days post vaccination, all mice were challenged with 10^2^ CCID_50_ (median cell culture infectious dose) of mouse-adapted HRTV delivered subcutaneously. The mouse-adapted HRTV (MA-HRTV) strain employed was derived from the MO-4 strain obtained from Dr. Robert Tesh (WRCEVA) [108]. The MA-HRTV stock (5.0 × 10^6^ CCID_50_/mL; 1 passage in Vero E6 cells, 5 passages of clarified liver homogenate in AG129 mice) used was prepared from a final passage in Vero E6 cells. The virus stock was diluted in sterile MEM and inoculated in injections of 200 μL. An additional group of two male and two female mice was sham infected with 200 μL MEM for use as a control. Following challenge, mice were weighed and evaluated daily for signs of infection.

We also evaluated the protective efficacy of passive serum transfer in AG129 mice. We pooled immune serum from the day 70 bleed of C57BL/6 mice vaccinated with rVSV-HRTV WT (p3) and rVSV-HRTV K1074A and evaluated the neutralizing titer of the pool via FRNT_50_. The approximate neutralizing antibody titer of the serum from rVSV-HRTV WT (p3) mice was 25, and the neutralizing titer from the mice vaccinated with rVSV-HRTV K1074A was approximately 253. Mice were injected subcutaneously with 10^2^ CCID_50_ mouse-adapted HRTV on day 0 and then were administered serum via IP injection on day 1. Four male and four female mice were included in each group. Mice were administered 120 μL of immune serum or 120 μL of serum diluted by 50% with sterile PBS, with one exception. Mice in the low dose rVSV-HRTV K1074A group received 110 μL of serum due to constraints on the volume of serum collected. Two additional groups were included as controls, and they were administered either 120 μL of PBS or 120 μL of non-immune serum from mice who had not been immunized against HRTV. Mice were weighed and evaluated for signs of infection for 21 days post challenge.

All mice were given approximately one week to acclimate to their cages and environment prior to initiation of experiments, and mice were maintained according to the Guide for Care and Use of Laboratory Animals. Vaccination experiments in C57BL/6 mice were performed under animal biosafety level 2 (ABSL-2) facilities at the University of Pennsylvania. Vaccination experiments in AG129 mice that included HRTV challenge were performed in ABSL-3 facilities at Utah State University. All animal work was approved by the University of Pennsylvania or Utah State University Institutional Animal Care and Use Committees (IACUC), and every effort was made to minimize pain and suffering of the mice.

### rVSV-HRTV Replication RT-qPCR

To quantify replication of rVSV-HRTV WT (p3), K1074A, and C621S+V753T+K1074A we evaluated viremia in vaccinated mice via quantitative PCR (qPCR). 50 μL of blood was collected from a subset of mice two days post vaccination, and 300 μL of TRIzol Reagent was added to each blood sample (Invitrogen, 15596026). RNA was extracted from each sample using the manufacturer’s protocol. cDNA was prepared from each RNA sample using the SuperScript III First-Strand Synthesis System according to the manufacturer’s protocol (Invitrogen, 18080-051). The final volume of cDNA was diluted 50% with sterile nuclease-free water and used for qPCR. For each reaction, we used 4.6 μL of PowerUp SYBR Green Master Mix (Applied Biosystems, A25742), 0.2 μL of each primer (10 μM), and 5 μL of diluted cDNA. Each cDNA sample was evaluated in triplicate using primers against VSV-N (Forward: 5’-GGAATACCCGGCAGATTACTT-3’; Reverse: 5’-GCCTTGGTAGACATATCCTCTTAG-3’) and mouse hypoxanthine-guanine phosphoribosyltransferase (mHPRT) (Forward: 5’-GTTGGATACAGGCCAGACTTTGTTG-3’; Reverse: 5’-GAGGGTAGGCTGGCCTATTGGCT-3’).

The qPCR assay was run in a 384-well plate on a QuantStudio 6 Flex Real-Time PCR System (Applied Biosystems). The following cycling conditions were used: Hold Stage: 50°C for 2 minutes, 95°C for 10 minutes; PCR Stage: 40 cycles of 95°C for 15 seconds, 60°C for 30 seconds, 72°C for 30 seconds; Melt Curve Stage: 95°C for 15 seconds, 60°C for 1 minute, 95°C for 15 seconds. Each assay was run using a VSV-N dilution series for positive controls and water for negative controls. The fold change of VSV-N (normalized to the mHPRT housekeeping gene) in vaccinated mice was compared to the sham control group of mice using the 2^-ΔΔCt^ method [116].

### Pseudovirus Neutralization Assays

Neutralizing antibodies produced in response to vaccination were evaluated via Focus Reduction Neutralization Tests (FRNTs). VSV pseudotypes with WT HRTV glycoproteins (VSVΔG-RFP(HRTV)) were produced as described above. Vero E6 cells were seeded in 100 μL at 2 × 10^4^ cells/well in a 96-well collagen coated plate. The next day, 2-fold serially diluted serum samples were mixed with VSVΔG-RFP(HRTV) pseudotype virus (100–200 focus forming units/well) and incubated for 1 hour at 37 °C. Also included in this mixture, to neutralize any potential VSV-G carryover virus, was 1E9F9, a mouse anti-VSV Indiana G, at a concentration of 600 ng/mL. The antibody-virus mixture was then used to replace the media on Vero E6 cells. Then, 30 hpi, the cells were washed with PBS and fixed with 4% paraformaldehyde before visualization on an S6 FluoroSpot Analyzer (CTL, Shaker Heights, OH, USA). Individual infected foci were enumerated, and the values compared to control wells without serum. The focus reduction neutralization titer 50% (FRNT_50_) was measured as the greatest serum dilution at which focus count was reduced by at least 50% relative to control cells that were infected with pseudotype virus in the absence of mouse serum. FRNT_50_ titers for each sample were measured in two to three technical replicates performed on separate days.

## Results

rVSV-SFTSV viruses are attenuated in cell culture, but serial passage of rVSV-SFTSV results in the spontaneous acquisition of mutations in the SFTSV glycoprotein that may improve viral replication [63, 64, 107]. To evaluate the impact of the mutations C617R, M749T, E982K, and K1071A on incorporation of the SFTSV glycoprotein into VSV particles, we developed CMV promoter plasmids encoding a codon-optimized coding region of the M segment of SFTSV and used mutagenesis to generate plasmids incorporating each mutation alone and in combination. These plasmids were used to produce VSVΔG(SFTSV) pseudotypes incorporating each mutant SFTSV glycoprotein (Figure 1A). VSVΔG pseudotypes do not encode a glycoprotein, and for these studies fluorescent RFP is inserted in the place of VSV-G. Though VSVΔG does not encode its own glycoprotein, budding VSVΔG particles can incorporate foreign viral glycoproteins expressed *in trans* through transfected or stably expressed genes. These virions enter cells using the biology of the incorporated glycoprotein they harbor but cannot produce infectious progeny in the target cell. Since these virions are only capable of a single round of replication, they are useful tools in studying various aspects of virion assembly without multiple rounds of replication yielding confounding results [111, 117]. Recovery of all SFTSV mutant pseudotypes indicates successful incorporation of each mutant glycoprotein onto VSV particles. We show that each substitution alone slightly increases the VSVΔG(SFTSV) titer, with E928K and M749T having the largest impact of single substitutions, resulting in an increase in titer of approximately one log. More importantly, synergism between substitutions is also evident, particularly with the inclusion of K1071A with E982K or M749T, resulting in a 95-fold increase over the WT titers (Figure 1A).

**Figure 1.**
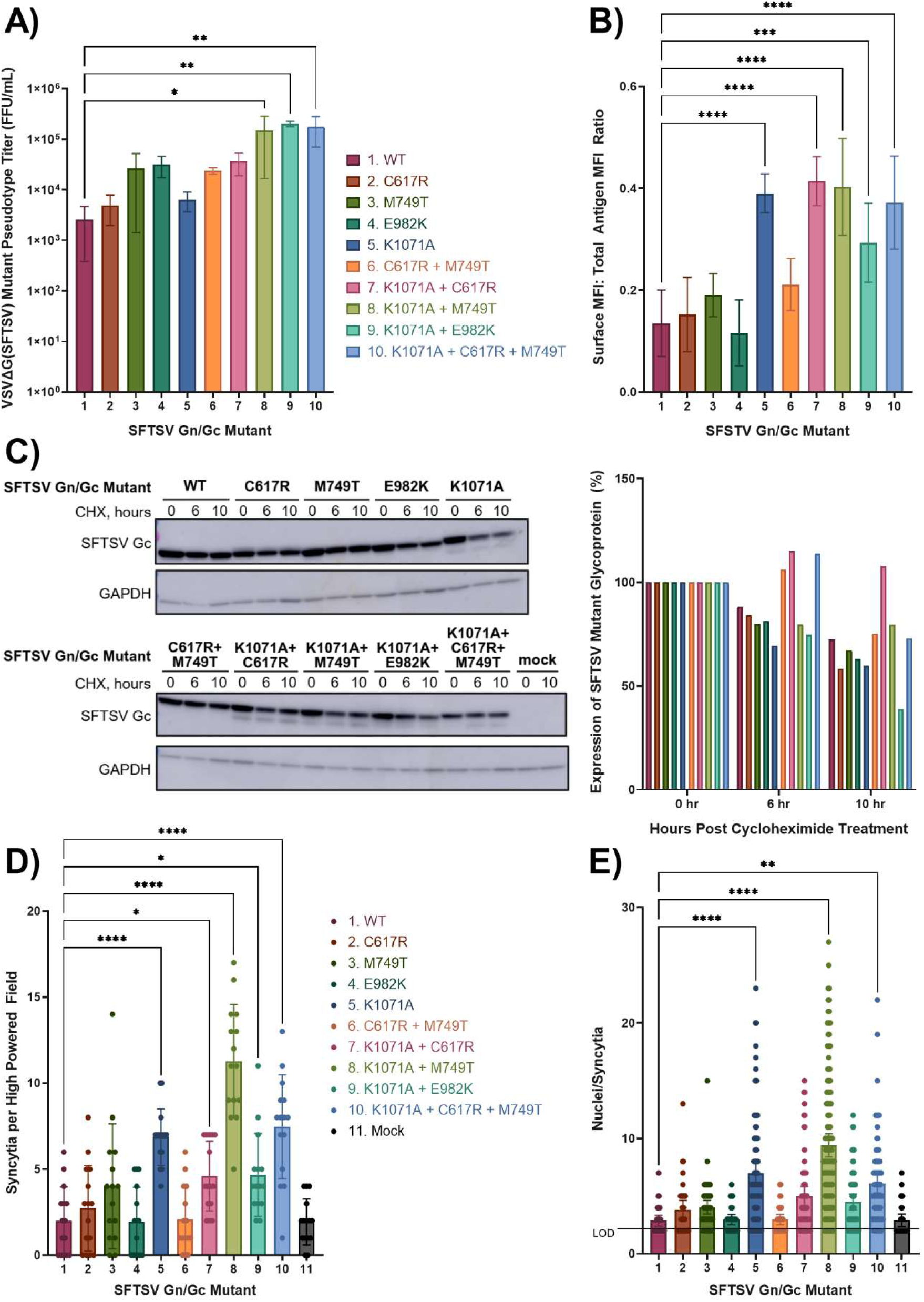
Mutations in SFTSV glycoprotein increase titers of VSVΔG pseudotypes via multiple mechanisms. **(A)** Titers of VSVΔG(SFTSV) mutant pseudotypes (FFU/mL) (Ordinary one-way ANOVA with Dunnett’s multiple comparisons test; *, *p <* 0.05; **; *p* < 0.01). **(B)** Surface expression of SFTSV mutant glycoprotein (ratio of MFI of surface SFTSV antigen to MFI of total antigen) (Ordinary one-way ANOVA with Dunnett’s multiple comparisons test; ***, *p <* 0.001; ****, *p* <0.0001). **(C)** Stability of SFTSV mutant glycoproteins treated with 50 μg/mL cycloheximide or DMSO as vehicle control and evaluated 6 hours and 10 hours post treatment. Protein expression in cells treated with cycloheximide was quantified via ImageJ and reported as a percentage of expression in DMSO treated cells. **(D)** Syncytia per high powered field and **(E)** nuclei per syncytia in cells transfected with SFTSV mutants and transiently treated with 50 mM MES buffered PBS at pH 5.5 (Ordinary one-way ANOVA with Dunnett’s multiple comparisons test; *, *p* <0.05; **, *p* <0.01; ****, *p* <0.0001; LOD: limit of detection, 2 nuclei).

As mentioned, VSVΔG pseudotypes are created when a budding VSV virion acquires a foreign viral glycoprotein, typically expressed on the plasma membrane due to the site of VSV assembly [111]. However, SFTSV glycoproteins are normally retained in the Golgi and ER-Golgi intermediate compartment (ERGIC) where SFTSV particles assemble and bud [118]. We hypothesized that the substitutions we observed may have a role in redistributing SFTSV glycoproteins to the surface of the cell, resulting in increased availability for incorporation into pseudotypes. To investigate this, we used flow cytometry to compare the expression of SFTSV glycoproteins on the surface of cells to SFTSV glycoproteins maintained in intracellular compartments (Figure 1B). This was done by staining surface proteins with mouse polyclonal serum reactive to SFTSV glycoproteins prior to permeabilization and labeling of internal proteins with rabbit polyclonal antibodies specific to SFTSV Gn and Gc. This allowed us to directly compare localization of the glycoproteins within individual cells. We found that K1071A has a large impact on surface expression, increasing the ratio of surface expressed protein to internally retained protein nearly four times over the WT glycoprotein. However, we did not see any further increase in surface expression with the addition of other substitutions. Given the phenotypes seen with the VSVΔG pseudotypes, it is surprising that no synergism is seen with the substitutions in terms of surface expression, and this suggests that there may be additional mechanisms responsible for the increased pseudotype titers.

To account for the effects of the substitutions that do not alter surface expression, we hypothesized that altered surface trafficking of the K1071A variant might result in exposure of SFTSV glycoproteins to low pH within the secretory pathway or to endocytosis and subsequent degradation from the cell membrane [119, 120]. The additional substitutions might serve to increase the stability of the K1071A protein, resulting in increased incorporation onto VSV particles and increased pseudotype titers. To assess SFTSV protein stability, we transfected cells with the mutant glycoproteins and treated the cells with DMSO or cycloheximide, which causes translation arrest. We harvested the cells 6 hours or 10 hours post treatment. Proteins were analyzed via Western blot using GAPDH as a loading control (Figure 1C). We found that glycoproteins with the K1071A substitution were much less stable, declining to approximately 52% protein expression at 10 hours post treatment compared to 79% protein expression with the WT glycoprotein. The addition of substitutions to K1071A, particularly K107A+C617R, seemed to increase the stability of the glycoprotein.

All the mutations identified are in the Gc component of the SFTSV glycoprotein, which is responsible for fusion and entry into cells [39–43]. Given this, we next hypothesized that the substitutions might affect pseudotype infectivity by impacting the fusogenicity of the glycoprotein. To evaluate this, we transfected cells with the mutant SFTSV glycoproteins and transiently treated the cells with an acidic buffer (pH 5.5) 48 hours post transfection. After low pH treatment, the cells were allowed to fuse for 4 hours before quantifying fusion by counting the number of syncytia in five high-powered fields per condition (Figure 1D) and the number of nuclei per syncytia (Figure 1E). As one might expect with an increase in surface expression, we saw that glycoproteins bearing the K1071A substitution resulted in overall increases in syncytia and the number of nuclei within. We saw further increases with K1071A+M749T, and this additional substitution resulted in the formation of an increased number and size of syncytia compared to K1071A alone. This indicates that increases in fusogenicity may play a role in the increased pseudotype titers. Overall, we find that substitutions in the SFTSV glycoprotein increase VSVΔG(SFTSV) pseudotype titers via multiple mechanisms.

The protein localization phenotype resulting from K1071A is striking, given that most bunyaviral glycoproteins are directed to internal compartments prior to budding [84–91]. Gn/Gc complexes encoded by most bunyaviruses localize to the Golgi when expressed separately from other viral factors [84, 86, 87, 90, 91, 93, 94, 121–123].The K1071A mutation is located in the cytoplasmic tail of the Gc component of the SFTSV glycoprotein, a region in the protein that has been implicated in ER targeting in other bunyaviruses [94]. More specifically, the K1071A substitution is located three amino acids from the carboxy terminus of the protein, otherwise referred to as the −3 position. This residue in the −3 position is accompanied by arginine, an additional positively charged amino acid, in the −5 position. This motif, RxKxx, resembles the canonical COPI binding signal KxKxx. COPI is a complex of several proteins that functions to recycle cargo from the Golgi to the ER and retain cargo within the ER, among other things. Cargo binding components of COPI typically interact with target proteins bearing dilysine motifs, either KKxx or KxKxx, that bind to α-COP or β’-COP respectively. Noncanonical β’-COP targeting sequences including KxHxx have been identified in the cytoplasmic tails of other viruses [96, 100, 104, 124–126]. We hypothesized that the K1071A substitution inhibits interaction with β’-COP via abrogation of a non-canonical RxKxx motif, resulting in increased surface expression.

To dissect the impact of the cytoplasmic tail mutation (K1071A) on Gc protein localization, we generated localization reporters consisting of SFTSV glycoprotein chimeras in which Gc ectodomain sequences were replaced with fluorescent reporters (mCherry or mNeonGreen). A549 cells were cotransfected with a plasmid encoding mNeonGreen appended to the WT Gc transmembrane domain and cytoplasmic tail (mNeonGreen-RxKxx) and a plasmid encoding mCherry appended to the transmembrane domain and cytoplasmic tail bearing the K1071A substitution (mCherry-RxAxx) (Figure 2A). In agreement with results from flow cytometry experiments (Figure 1B), mNeonGreen-RxKxx exhibited an ER distribution pattern with undetectable levels at the cell surface. In contrast, mCherry-RxAxx was clearly detectable at the cell surface (Figure 2A). This suggests that the RxKxx motif alone is sufficient to distribute proteins to the ER, regardless of the identity of the ectodomain, and that mutation of the −3 lysine abrogates the ER retention. To quantify the phenotype exhibited in Figure 2A, we transfected cells with mCherry-RxKxx, mCherry-RxAxx, or cytosolic mCherry to compare the surface expression of mCherry via flow cytometry. Non-permeabilized cells were stained with an antibody specific to mCherry and labeled with Alexa Fluor 488 to differentiate surface antigen from the intracellular mCherry signal. The percentage of total mCherry protein that was expressed on the surface of cells is reported in Figure 2B. As expected from the IF results, mCherry-RxKxx was predominantly expressed inside the cell, and only a small percentage (less than 30%) of the total protein was expressed at the cell surface. In accordance with previous results, mCherry-RxAxx exhibited significantly more cell surface expression with greater than 64% of proteins trafficking to the cell surface. This represents a greater than 2-fold increase in cell surface expression when comparing a construct with the Gc WT transmembrane domain and cytoplasmic tail to one with the −3 lysine to alanine mutation (Figure 2B).

**Figure 2.**
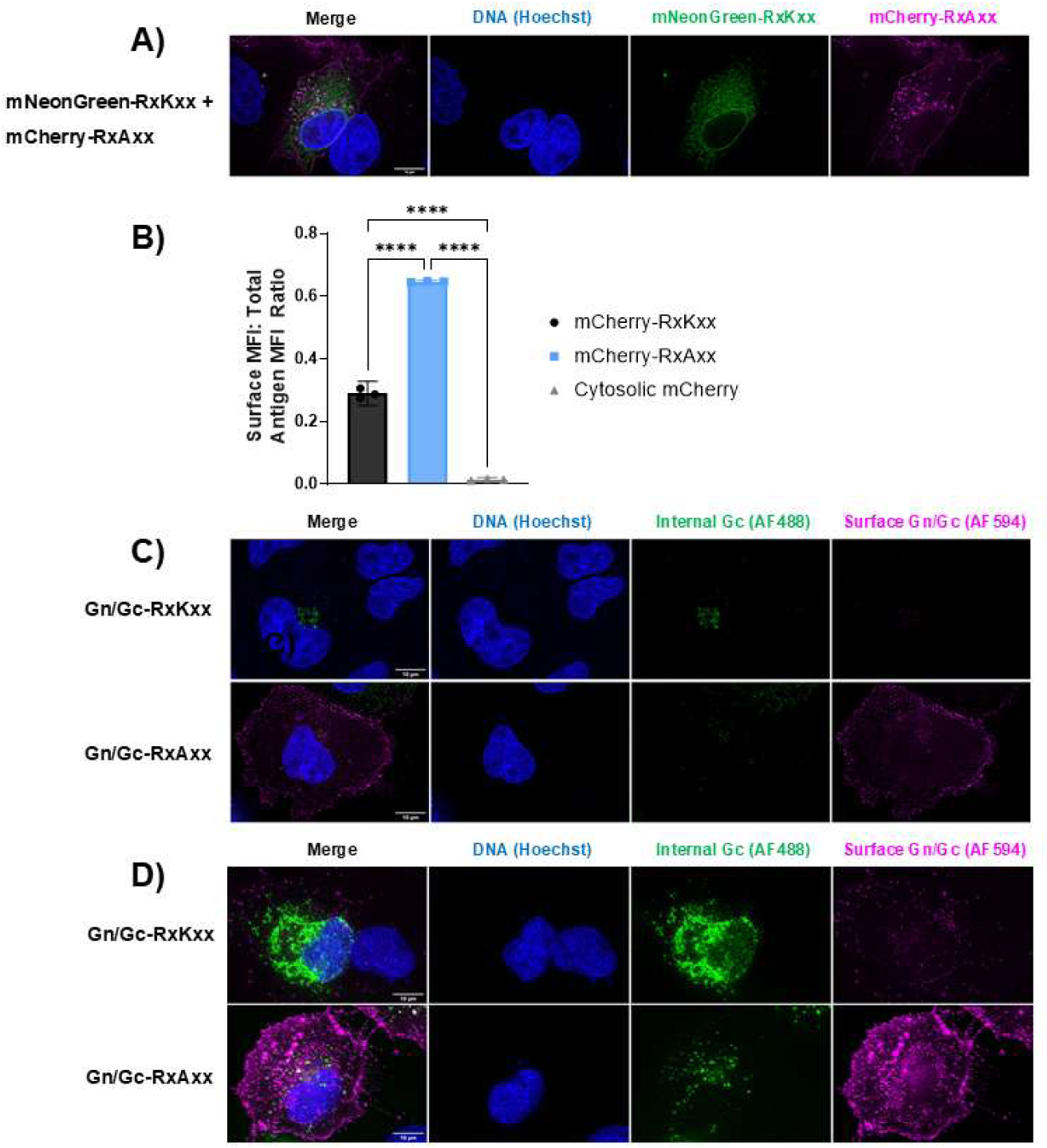
K1071A substitution redistributes SFTSV Gn and Gc from intracellular compartments to the cell surface. **(A)** Fluorescence microscopy of cells co-expressing mNeonGreen-RxKxx and mCherry-RxAxx from separate plasmids. **(B)** Surface expression of mCherry (ratio of MFI of surface mCherry to MFI of total mCherry antigen) (Ordinary one-way ANOVA with Tukey’s multiple comparisons test; ****, *p* <0.0001). Data is representative of three independent experiments. **(C)** Immunofluorescence microscopy of internal (permeabilized) and surface antigen in transfected A549 cells expressing wild type (RxKxx) or K1071A mutant (RxAxx) SFTSV Gn/Gc. **(D)** Immunofluorescence microscopy of internal (permeabilized) and surface antigen in transfected U2OS cells expressing wild type (RxKxx) or K1071A mutant (RxAxx) SFTSV Gn/Gc. Images in **(A,C, and D)** were collected at 100x under oil immersion, and scale bars are 10 μm in length. Images are of representative equatorial slices.

Evidence that Gn dictates Golgi targeting include the observations that Gn expressed in isolation of Gc localizes to the Golgi, and recruitment of Gc from the ER to the Golgi requires co-expression with Gn. The contribution made by Gc to the overall distribution of Gn/Gc complexes has been less well-studied and remains unclear [93, 94]. To study the effect of the K1071A substitution on the localization of the Gn/Gc complex, A549 cells were transfected with expression plasmids encoding SFTSV Gn/Gc with the WT tail sequence (Gn/Gc-RxKxx) or Gn/Gc with the K1071A substitution (Gn/Gc-RxAxx). Similar to the methods used for Figure 1B, external proteins were labeled with mouse polyclonal serum reactive to SFTSV glycoproteins and internal proteins were labeled with rabbit polyclonal antibodies specific for SFTSV Gn and Gc, and cells were imaged by immunofluorescence microscopy. Surface signal from cells expressing Gn/Gc-RxKxx was low or absent in most antigen-positive cells (Figure 2C). Internal signal in most cells was concentrated in a perinuclear region that was likely the Golgi. In contrast, surface signal intensity of cells transfected with Gn/Gc-RxAxx was significantly greater, and full cell boundaries were visible. Gn/Gc-RxAxx also distributed differently within intracellular compartments. Given the decreased stability of proteins with the K1071A substitution (Figure 1C), it is possible that the proteins are endocytosed from the cell surface and targeted for lysosomal degradation, which may partially explain the altered intracellular distribution. Expression in a different cell line, U2OS, displayed similar staining patterns for Gn/Gc-RxKxx and Gn/Gc-RxAxx to those observed in A549 cells, however the surface localization of the RxAxx mutant is more obvious (Figure 2D). Taken together, these data suggest that the K1071A substitution redistributes Gn/Gc complexes from intracellular compartments to the cell surface.

Given the significant impact of the K1071A substitution on protein trafficking and surface expression of the SFTSV glycoprotein and our hypothesis that the K1071A substitution altered a potential noncanonical COPI binding site, we wanted to interrogate the mechanism responsible for that phenotype more thoroughly. We used transient SFTSV Gn/Gc expression to address via immunofluorescence microscopy whether the viral glycoproteins colocalize with endogenous β’-COP. A549 cells were transfected with the WT plasmid (Gn/Gc-RxKxx) or the K1071A plasmid (Gn/Gc-RxAxx) and stained with antibodies against SFTSV Gn/Gc and β’-COP. As a component of the coatomer complex I (COPI), which is involved in retrograde trafficking within the Golgi cisternae and between the Golgi and ER, β’-COP is localized primarily to the Golgi in mammalian cells [127, 128]. Treatment of cells with the microtubule depolymerizing agent nocodazole disrupts Golgi structure, relocalizes Golgi resident proteins into dispersed cytoplasmic vesicles, and allows improved spatial resolution between the Golgi and adjacent structures [129]. In cells treated with vehicle, β’-COP staining was most intense in a perinuclear region consistent with the Golgi (Figure 3A). Gn/Gc-RxKxx and Gn/Gc-RxAxx signal overlapped with that of β’-COP, but Gn/Gc-RxAxx appeared more dispersed through the cytosol. Similar results were seen in cells transiently expressing Gn/Gc-RxKxx or Gn/Gc-RxAxx that were treated with nocodazole before fixation and staining for viral antigen and β’-COP (Figure 3C). The fractional overlap between Gn/Gc-RxAxx and β’-COP was reduced compared to that between Gn/Gc-RxKxx and β’-COP under both mock (Figure 3B) and nocodazole-treated conditions (Figure 3D). These results suggest that the fraction of Gn/Gc-RxAxx retained in the Golgi is reduced compared to Gn/Gc-RxKxx.

**Figure 3.**
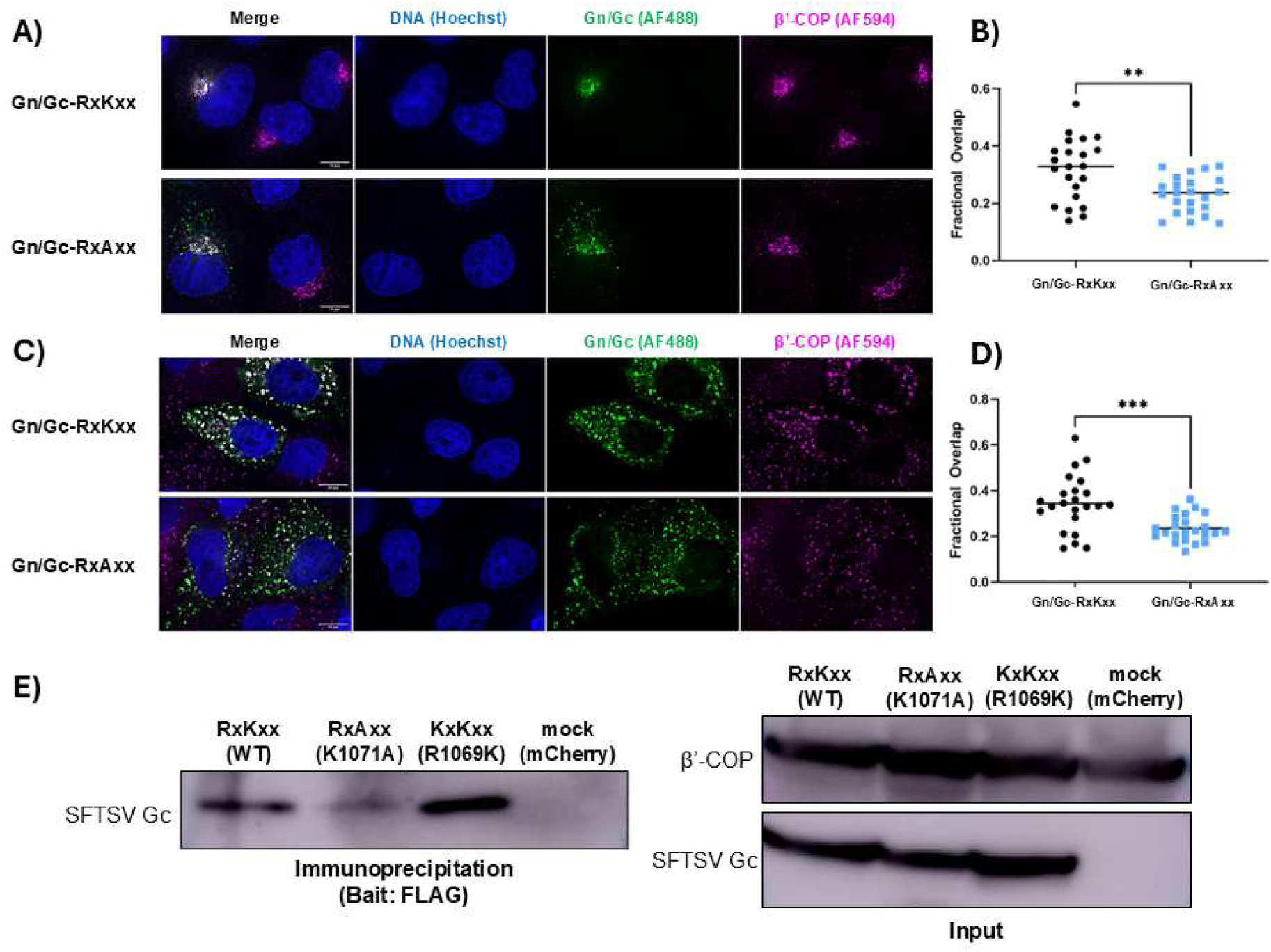
K1071A mutation increases surface expression of SFTSV glycoproteins via inhibition of COPI binding. **(A)** Immunofluorescence microscopy of vehicle (DMSO)-treated A549 cells transiently expressing SFTSV Gn/Gc-RxKxx or Gn/Gc-RxAxx. Cells were stained for the Golgi marker β’-COP. Images were acquired at 100x magnification. Images are representative equatorial slices. Scale bars are 10 μm in length. **(B)** Fractional overlap of Gn/Gc-RxKxx or Gn/Gc-RxAxx and β’-COP. Dots represent single cells (n = 22 cells/condition) pooled from two independent experiments (Student’s t-test; **, *p* < 0.01). **(C)** Immunofluorescence microscopy of A549 cells transfected to express SFTSV Gn/Gc-RxKxx or Gn/Gc-RxAxx and pretreated for 3 hours with 33 μM nocodazole then stained for endogenous β’-COP. Images were acquired at 100x magnification. Images are of representative equatorial slices. Scale bars are 10 μm in length. **(D)** Fractional overlap of Gn/Gc-RxKxx or Gn/Gc-RxAxx with β’-COP. Each dot represents an individual cell with n = 23 cells/condition collected from two independent experiments (Student’s t-test; ***, *p* < 0.001). **(E)** Immunoprecipitation of SFTSV Gn/Gc-RxKxx, -RxAxx, or -KxKxx (positive control) from 293T cells cotransfected with SFTSV Gn/Gc-RxKxx, -RxAxx, or –KxKxx and FLAG-tagged β’-COP. Protein was bound to magnetic beads conjugated to an anti-FLAG antibody, washed thoroughly with cell lysis buffer, and eluted via boiling. Proteins recovered from IP were evaluated via SDS-PAGE and Western blot using an antibody against SFTSV Gc and an antibody against β’-COP. Data are representative of four independent experiments

Finally, to directly address the hypothesis that the K1071A substitution increases surface expression via inhibition of binding to β’-COP, we evaluated binding of SFTSV Gn/Gc-RxKxx or Gn/Gc-RxAxx to β’-COP via an immunoprecipitation assay. We cotransfected HEK293T cells with Gn/Gc-RxKxx or Gn/Gc-RxAxx expression plasmids along with a commercially available β’-COP plasmid tagged with FLAG at the carboxy terminus. This commercially available β’-COP plasmid was chosen based on reports that a C-terminal tagged β’-COP is capable of binding cargo [130, 131]. An expression plasmid with a R1069K substitution (Gn/Gc-KxKxx) was used as a positive control, as we expected that a canonical COPI binding motif in the cytoplasmic tail would bind to β’-COP more efficiently. A plasmid expressing mCherry was used as a negative control. We harvested cell lysate 48 hours post transfection and precipitated proteins using anti-FLAG beads. The beads were thoroughly washed using cell lysis buffer, and bound protein was eluted via boiling. We found that Gn/Gc carrying the wild type RxKxx motif co-precipitates with β’-COP. As anticipated, we found that Gn/Gc with a canonical binding motif, KxKxx, co-precipitates with β’-COP with the highest efficiency of the proteins tested. Very little immunoprecipitation of the mutant Gn/Gc-RxAxx protein was seen, which supports our hypothesis that the K1071A substitution inhibits binding to β’-COP (Figure 3E).

Next, we wanted to understand the effects of the mutations within the glycoprotein on replication-competent rVSV-SFTSV. Using published VSV launch strategies [109], we attempted to launch rVSV-SFTSV viruses bearing all the substitution combinations seen in Figure 1. Despite numerous attempts, we were only able to launch a subset of these viruses: rVSV-SFTSV K1071A, rVSV-SFTSV K1071A+C617R, rVSV-SFTSV K1071A+M749T, and rVSV-SFTSV M749T. We were unable to launch rVSV-SFTSV bearing the WT glycoprotein that contains a non-canonical COPI binding motif. Our lab first launched rVSV-SFTSV in 2013 for use in identifying SFTSV entry factors [44]. Sequencing of the earliest stocks of this virus revealed that the K1071 residue was mutated, suggesting that this substitution may have been required for launch or effective replication after launch.

Following launch of rVSV-SFTSV K1071A, rVSV-SFTSV K1071A+C617R, rVSV-SFTSV K1071A+M749T, and rVSV-SFTSV M749T, we passaged each virus five times and sequenced the SFTSV glycoprotein in each passage to determine if the mutations were genetically stable. We found that rVSV-SFTSV K1071A was stable across five passages, but each other viral mutant acquired additional mutations. Surprisingly, rVSV-SFTSV K1071A+C617R acquired the E982K substitution in the first passage after launch. rVSV-SFTSV K1071A+M749T acquired a substitution in the transmembrane domain of Gn, L462P, by passage 3. rVSV-SFTSV M749T was the least stable of the viruses, picking up numerous mutations that changed between passages (Figure 4A). For consistency and brevity, we will continue to refer to each virus with the mutations that were present at launch, not the mutations that arose spontaneously over passage.

**Figure 4.**
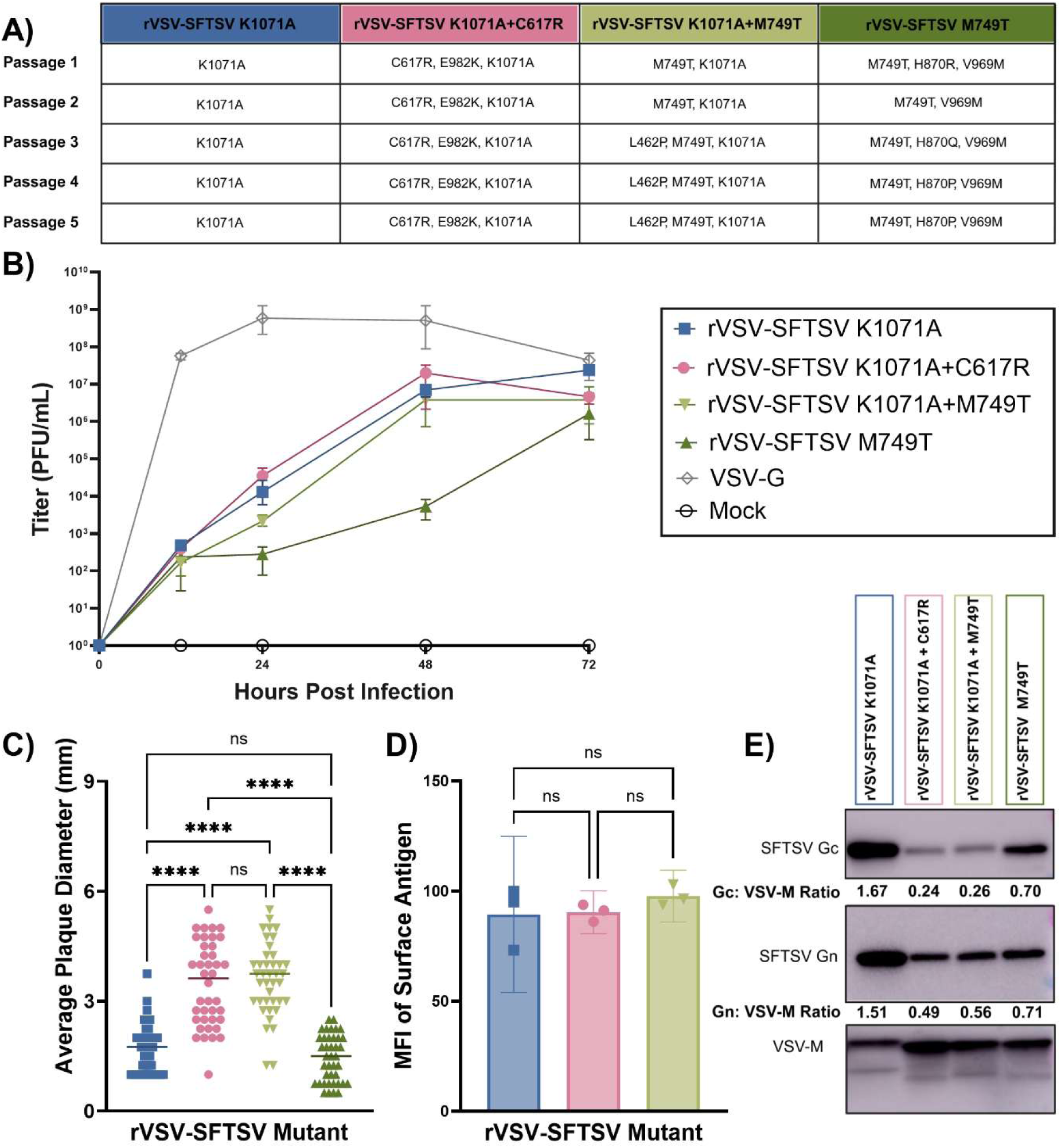
K1071A mutation enhances replication of rVSV-SFTSV and improves incorporation of SFTSV onto VSV virions. **(A)** Five passages of each rVSV-SFTSV mutant were sequenced via nanopore sequencing to assess genetic stability of the SFTSV glycoprotein. Nonsynonymous mutations are reported. **(B)** Replication kinetics of rVSV-SFTSV mutants compared to VSV-G. Vero E6 cells were infected at an MOI of 0.001, and samples of supernatant were collected at 12-, 24-, 48-, and 72-hours post infection. Viral titers were evaluated via plaque assays on Vero E6 cells. **(C)** Plaques from each rVSV-SFTSV mutant were measured with a standard ruler and rounded to the nearest quarter of a mm (Ordinary one-way ANOVA with Tukey’s multiple comparisons test; ****, *p* < 0.0001). **(D)** Surface expression of mutant glycoproteins in A549 cells infected with rVSV-SFTSV mutants at an MOI of 2. Cells were fixed and stained for flow cytometry 24 hours post infection (Ordinary one-way ANOVA with Tukey’s multiple comparisons test; no significant differences found). These data are representative of three independent experiments. **(E)** Incorporation of mutant glycoproteins onto rVSV particles. 15 cm plates of Vero E6 cells were infected with each rVSV-SFTSV mutant at an MOI of 0.001. 48 hours post infection, supernatant was harvested, and virions were purified via ultracentrifugation through a sucrose cushion at 28,000xg for 2 hours. Virions were lysed using Triton X-100, and protein samples were evaluated via SDS-Page and Western blot using antibodies specific to SFTSV Gn, Gc, and VSV-M. Ratios of Gn or Gc to VSV-M were calculated using ImageJ. These data are representative of two independent experiments.

To determine if the mutations impacted the replication of rVSV-SFTSV, we infected Vero E6 cells with the rVSV-SFTSV mutants at a low MOI and collected virus-containing supernatant at 12-, 24-, 48-, and 72-hours post infection. We used standard plaque assays to evaluate the titer over the course of the replication experiment, and titers were compared to VSV with the native VSV-G glycoprotein. All the rVSV-SFTSV viruses replicated to high titers that would be appropriate for large scale experiments, but they were still highly attenuated compared to VSV. All rVSV-SFTSV variants bearing the K1071A substitution reached a maximum titer of approximately 10^7^ PFU/mL, but rVSV-SFTSV M749T reached a peak titer of 10^6^ PFU/mL. All the viruses bearing the K1071A substitution replicated significantly more rapidly than the virus with only the M749T substitution, but we were surprised to find that there was no significant difference in replication between rVSV-SFTSV K1071A, rVSV-SFTSV K1071A+C617R, and rVSV-SFTSV K1071A+M749T (Figure 4B).

The plaque assays used to calculate the titers also revealed an interesting phenotype in plaque sizes in Vero E6 cells. All rVSV-SFTSV viruses produced distinguishable plaques 96 hours post infection. However, there is a significant difference in the size of plaques at this timepoint. rVSV-SFTSV K1071A plaques are small, with an average diameter of 1.7 mm. The plaques for the K1071A+C617R and K1071A+M749T viruses are significantly larger and have an average diameter of 3.5 and 3.6 mm, respectively (Figure 4C). This may reflect increased cell-cell spread, potentially resulting from increased fusogenicity of the glycoproteins. Supporting this idea, we previously demonstrated increased fusogenicity of the K1071A+M749T glycoprotein mutant (Figure 1D-E). The recombinant virus carrying the M749T variant produced small plaques that were comparable to rVSV-SFTSV K1071A (Figure 4C).

Although our flow cytometry data exploring the surface expression of the mutant glycoprotein plasmids suggested that there was not a significant difference in surface localization in glycoproteins bearing the K1071A substitution (Figure 1B), we wanted to rule this out as an explanation for the differences in plaque sizes. We infected A549 cells at an MOI of 2 and evaluated surface glycoprotein expression in infected cells by flow cytometry. We excluded rVSV-SFTSV M749T from this analysis due to its slow replication, which prevented analysis of surface expression at the same timepoint as the other viruses. As expected, we found that there was no significant difference in surface expression between the K1071A, K1071A+C617R, and K1071A+M749T recombinant viruses (Figure 4D). This suggests that the differences in plaque sizes result from a mechanism independent of surface expression.

Finally, we evaluated incorporation of the glycoproteins into purified rVSV virions via a Western blot comparing expression of SFTSV Gn and Gc to VSV-M. We found that rVSV-SFTSV K1071A Gn and Gc were incorporated much more efficiently than any of the other mutants, with an expression of 1.51 and 1.67 times VSV-M, respectively. Unexpectedly, Gc of rVSV-SFTSV K1071A+C617R and rVSV-SFTSV K1071A+M749T were not efficiently incorporated, with an expression of approximately 0.25 times VSV-M for both viruses (Figure 4E).

We next sought to determine whether the mutations identified in SFTSV could be relevant to other viruses in the *Bandavirus* genus. Heartland virus (HRTV) is a closely related bandavirus with a high case fatality rate, and no vaccines have yet been developed to prevent infection [31]. We wanted to develop an rVSV-HRTV vaccine for pre-clinical testing, and we hypothesized that the SFTSV substitutions might improve replication of the virus, resulting in improved ease of production and enhanced immunogenicity. Notably, HRTV carries a non-canonical COPI binding motif RxKxx, identical to the one in SFTSV.

Based on our initial data from the VSVΔG(SFTSV) pseudotypes (Figure 1A) and surface expression (Figure 1B), we decided to focus on the K1071A substitution and the combination of K1071A+C617R+M749T substitutions. Using sequence homology, we built these substitutions into the cognate positions of the HRTV glycoprotein. For HRTV, this corresponded to K1074, C621, and V753. For K1074 and V753, we made the same substitutions used in SFTSV: K1074A and V753T. However, we mutated C621 to a serine based on published data that showed a C621S substitution occurred spontaneously in HRTV passaged in mice [108]. Since this substitution seemed to improve replication of HRTV in animals, we hypothesized that it would also be beneficial in rVSV-HRTV.

Using the same rVSV launch protocol as described above, we attempted to launch rVSV-HRTV encoding the WT, K1074A, and C621S+V753T+K1074A variants. All these viruses were successfully rescued, which was a surprise to us given the difficulty we had in recovering rVSV-SFTSV WT. We again sequenced the viruses to determine if the genome was stable over multiple passages. We found that rVSV-HRTV WT was stable to passage 2 but acquired two additional mutations by passage 3, causing the changes H572R and N874K. rVSV-HRTV C621S+V753T+K1074A acquired one additional mutation by passage 2, a V1052M substitution. In contrast, rVSV-HRTV K1074A was stable through passage 5 (Figure 5A).

**Figure 5.**
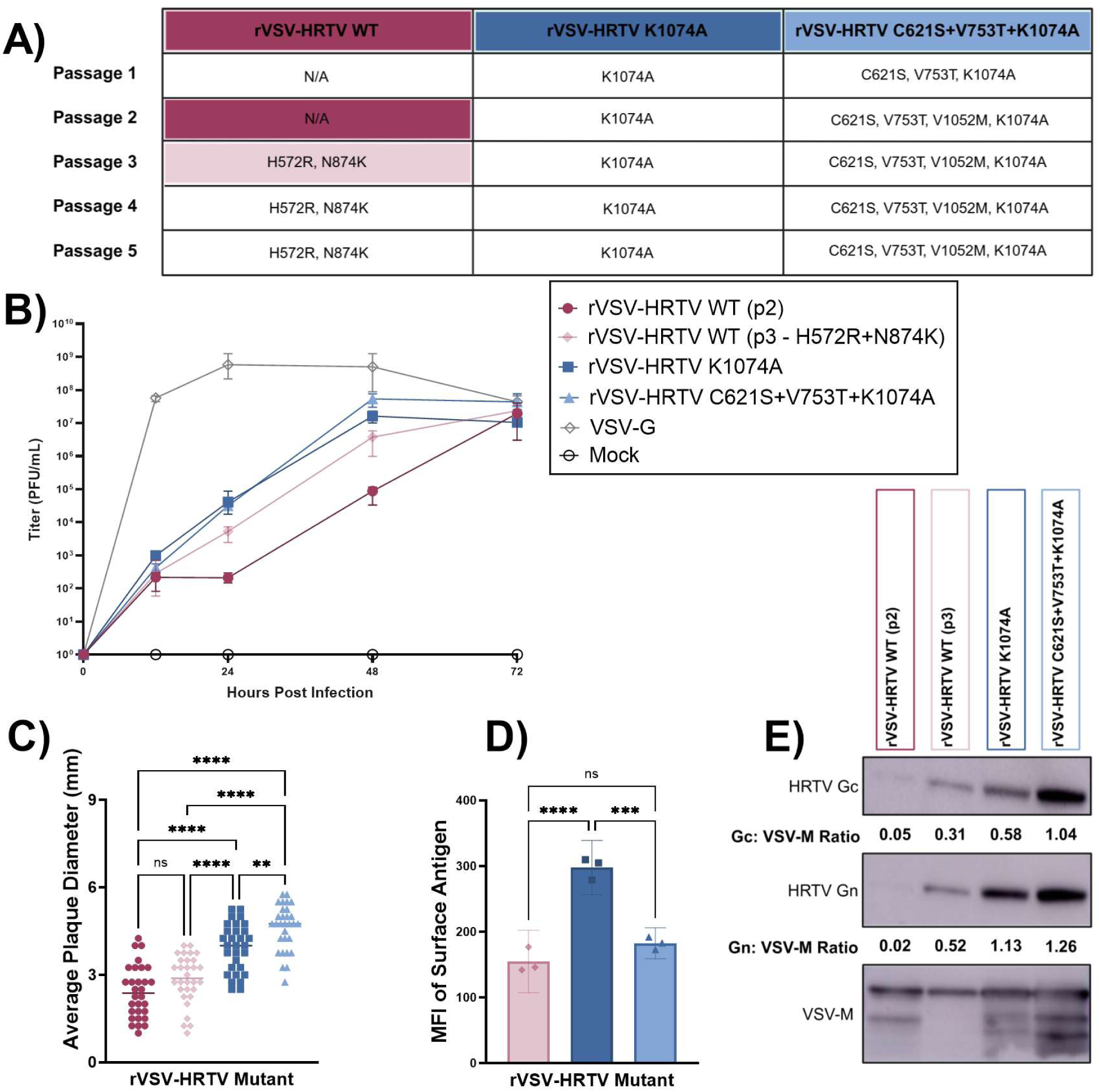
K1074A mutation is also effective in rVSV-HRTV, leading to enhanced replication and glycoprotein expression on the surface of infected cells. **(A)** Five passages of each rVSV-HRTV mutant were sequenced via nanopore sequencing to assess genetic stability of the HRTV glycoprotein. Nonsynonymous mutations are reported. **(B)** Replication kinetics of rVSV-HRTV mutants compared to VSV-G. Vero E6 cells were infected at an MOI of 0.001, and samples of supernatant were collected at 12-, 24-, 48-, and 72-hours post infection. Viral titers were evaluated via plaque assays on Vero E6 cells. **(C)** Plaques from each rVSV-HRTV mutant were measured with a standard ruler and rounded to the nearest quarter of a mm (Ordinary one-way ANOVA with Tukey’s multiple comparisons test; **, *p* < 0.01; ****, *p* < 0.0001). **(D)** Surface expression of mutant glycoproteins in A549 cells infected with rVSV-HRTV mutants at an MOI of 2. Cells were fixed and stained for flow cytometry 24 hours post infection (Ordinary one–way ANOVA with Tukey’s multiple comparisons test; ***, *p <* 0.001; ****, *p* < 0.0001). These data are representative of three independent experiments. **(E)** Incorporation of mutant glycoproteins onto rVSV particles. 15 cm plates of Vero E6 cells were infected with each rVSV-HRTV mutant at an MOI of 0.001. 48 hours post infection, supernatant was harvested, and virions were purified via ultracentrifugation through a sucrose cushion at 28,000×g for 2 hours. Virions were lysed using Triton X-100, and protein samples were evaluated via SDS-Page and Western blot using antibodies specific to VSV-M and SFTSV antibodies that cross react with HRTV Gn and Gc. Ratios of Gn or Gc to VSV-M were calculated using ImageJ. These data are representative of two independent experiments.

Following launch of the viruses, we did a time course experiment using the same methods as those used for Figure 4B. For comparison, we included the second passage of rVSV-HRTV WT because sequencing showed that this passage had not acquired any additional mutations (Figure 5A). Titers at each timepoint were again compared to those of VSV-G. We saw very similar replication kinetics between rVSV-SFTSV K1071A and rVSV-HRTV K1074A, with similar titers at each timepoint and peak titers reaching approximately 10^7^ PFU/mL. Again, we did not see a significant difference between rVSV-HRTV K1074A and rVSV-HRTV C621S+V753T+K1074A, suggesting that the K1074A substitution is sufficient to improve rVSV-HRTV replication, as was the case with rVSV-SFTSV. rVSV-HRTV WT (p2) replicated significantly slower than the viruses with the K1074A substitution, resulting in a difference in titer of approximately 2.5 logs at 24 hours post infection and 2 logs at 48 hours post infection. rVSV-HRTV WT (p3), which had acquired additional changes, grew with an intermediate phenotype, replicating faster than WT (p2) but slower than the viruses with the K1074A substitution (Figure 5B). This suggests that there are multiple mutation pathways that the virus can take to improve replication *in vitro*.

We measured the sizes of plaques formed by each rVSV-HRTV mutant, and again rVSV-HRTV WT (p2) was included as a control. No significant difference in plaque sizes was seen between passage 2 and 3 of rVSV-HRTV WT, suggesting that the two additional mutations acquired spontaneously do not appear to impact the cell-cell spread of the virus. Plaques for rVSV-HRTV K1074A were significantly larger than either of the rVSV-HRTV WT passages, but rVSV-HRTV C621S+V753T+K1074A formed the largest plaques of any of the viruses (Figure 5C). This may reflect that the additional mutations aid in fusogenicity and cell-cell spread, as was suggested by the rVSV-SFTSV results.

Next, we evaluated the cell surface expression of glycoproteins in cells infected with the rVSV-HRTV mutants by flow cytometry. We excluded rVSV-HRTV WT (p2) due to its slower replication, which prevented analysis at the same timepoint as the other viruses. As expected, we found a significant difference in surface expression between rVSV-HRTV WT (p3) and rVSV-HRTV K1074A, which indicates that the substitution of the lysine residue at the −3 position is responsible for redistribution of glycoproteins to the cell surface in HRTV as well as SFTSV. To our surprise, we found that the surface expression of rVSV-HRTV C621S+V753T+K1074A was significantly lower than rVSV-HRTV K1074A and was comparable to rVSV-HRTV WT (Figure 5D). It is unclear why this would be the case, especially given the similarity in replication kinetics between rVSV-HRTV K1074A and rVSV-HRTV C621S+V753T+K1074A.

We also evaluated incorporation of HRTV Gn and Gc into VSV virions. Incorporation of Gn and Gc from HRTV WT (p2) was extremely inefficient, and these proteins were barely detectable in purified virions. The substitutions that arise spontaneously in rVSV-HRTV WT by passage 3, H572R and N874K, improve incorporation of the glycoproteins relative to passage 2. The K1074A substitution dramatically improves glycoprotein incorporation relative to rVSV-HRTV WT (p3), but the combination of C621S+V753T+K1074A substitutions results in the most efficient incorporation of HRTV glycoproteins into virions (Figure 5E).

To evaluate replication of the rVSV-HRTV vaccine candidates in animal models, we injected groups of 6 male and 6 female AG129 mice with a low dose (10^2^ PFU) of each vaccine candidate via the intraperitoneal route. AG129 mice are deficient in interferon α, β, and γ receptors, which makes them highly susceptible to viral infections and allows observation of viral replication *in vivo* [132, 133]. These mice also serve as the established HRTV challenge model [108] allowing evaluation of the vaccine potential for these rVSV-HRTV candidates. In response to infection with rVSV-HRTV WT (p3, which carries two spontaneous changes), the mice lost a moderate amount of weight starting at day 3 post injection, but they all recovered by day 7 (Figure 6A). This suggests that the virus replicates well initially in these mice before they are able to control replication. Infection with rVSV-HRTV C621S+V753T+K1074A had similar results, but this virus was better tolerated by the mice, and they only lost a small amount of weight (Figure 6A). In contrast, rVSV-HRTV K1074A immunization resulted in weight loss in the mice similar to that of the rVSV-HRTV WT group (Figure 6A), but most (10 of the 12 mice) succumbed within 5-7 days post injection (Figure 6B). This might suggest that rVSV-HRTV K1074A replicates more efficiently *in vivo* than the other mutants, resulting in mortality in severely immunocompromised mice. To directly address in vivo replication, we evaluated rVSV viremia in vaccinated mice two days post vaccination. We used qPCR to quantify levels of VSV-N RNA in the blood of vaccinated mice relative to the mouse housekeeping gene hypoxanthine guanine phosphoribosyltransferase (mHPRT). The average fold change of VSV-N in vaccinated mice relative to the sham infected control group was quantified via the 2^-ΔΔCt^ method [116]. We found that mice vaccinated with rVSV-HRTV WT or rVSV-HRTV C621S+V753T+K1074A did not exhibit detectable replication on day 2 post vaccination compared to the control group, with an average fold change of 1.6 or 0.8, respectively. In contrast, we observed significant replication of rVSV-HRTV K1074A two days post vaccination. The average fold change in VSV-N RNA was 103.6 times higher than the sham control group, confirming replication of this rVSV in AG129 mice.

**Figure 6.**
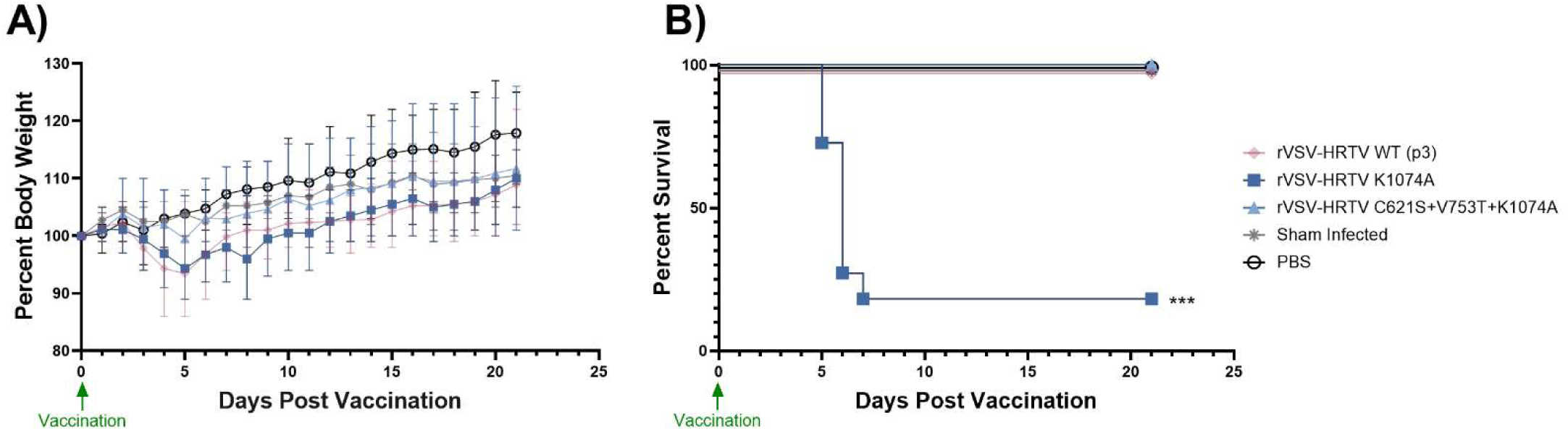
**rVSV-HRTV K1074A exhibits enhanced replication *in vivo.*** Six male and six female AG129 mice were IP injected with 102 PFU of rVSV-HRTV vaccine candidates. **(A)** Weight loss (mean and range) and **(B)** survival curves are shown for 21 days following injection. Statistical significance of survival was evaluated via Mantel-Cox logrank test compared to mice vaccinated with PBS placebo (***, *p* <0.0001).

Our ultimate goal of this study was to identify mutations in rVSV-SFTSV and rVSV-HRTV that would be genetically stable, improve our ability to produce high levels of the viruses for clinical studies, and contribute to a more immunogenic vaccine candidate. We previously published work on rVSV-SFTSV, where we showed that a single dose of an rVSV-SFTSV vaccine with the K1071E and E982K substitutions was protective against lethal SFTSV challenge and also provided cross-protection against HRTV [64, 112]. To determine if substitutions that affect HRTV glycoprotein function *in vitro* and *in vivo* replication would also improve the efficacy of an rVSV-HRTV vaccine candidate, we characterized our rVSV-HRTV mutants *in vivo* and evaluated safety, immunogenicity, and protection against lethal challenge.

First, we vaccinated groups of 8-week-old C57BL/6 mice with two doses (10^6^ PFU each) of each virus via intraperitoneal injections (IP). For this experiment, 5 male and 5 female mice were included in each group. Mice were given the prime vaccination dose at day 0 and boosted at day 28. Mice were bled via the facial vein prior to vaccination (day 0), and at days 14, 26, 42, 56, and 70 post prime vaccination and serum was isolated and stored for analysis. We employed focus reduction neutralization tests to define the dilution at which 50% of a VSVΔG(HRTV) pseudovirus was neutralized by the serum (FRNT_50_). We found that neutralizing antibody titers in these immunocompetent C57BL/6 mice were relatively low across all groups prior to the boost. However, rVSV-HRTV K1074A vaccination induced significantly higher titers than rVSV-HRTV WT (p3) and rVSV-HRTV C621S+V753T+K1074A at day 26 post vaccination. Following the boost vaccination at day 28, we saw an increase in neutralizing titers in all vaccine groups. At each timepoint following the boost, rVSV-HRTV K1074A outperformed rVSV-HRTV WT (p3) by inducing significantly higher neutralizing antibody titers. Post-boost neutralizing antibody titers were not significantly different between rVSV-HRTV K1074A and rVSV-HRTV C621S+V753T+K1074A, however the trend shows higher neutralizing titers on average when mice were vaccinated with rVSV-HRTV K1074A. This suggests that rVSV-HRTV with the single K1074A substitution is superior to the other two mutants tested, which may be a result of the increased surface expression of glycoproteins with this substitution (Figure 5D) or increased replication *in vivo* (Figure 6A-B). However, it’s also important to note that we saw high variability in neutralizing antibody titers in all vaccine groups, which may suggest variable levels of viral replication between immunocompetent individuals. Nevertheless, every animal vaccinated with rVSV-HRTV K1074A seroconverted, which was not the case for animals vaccinated with rVSV-HRTV WT (p3) and rVSV-HRTV C621S+V753T+K1074A (Figure 7A).

**Figure 7.**
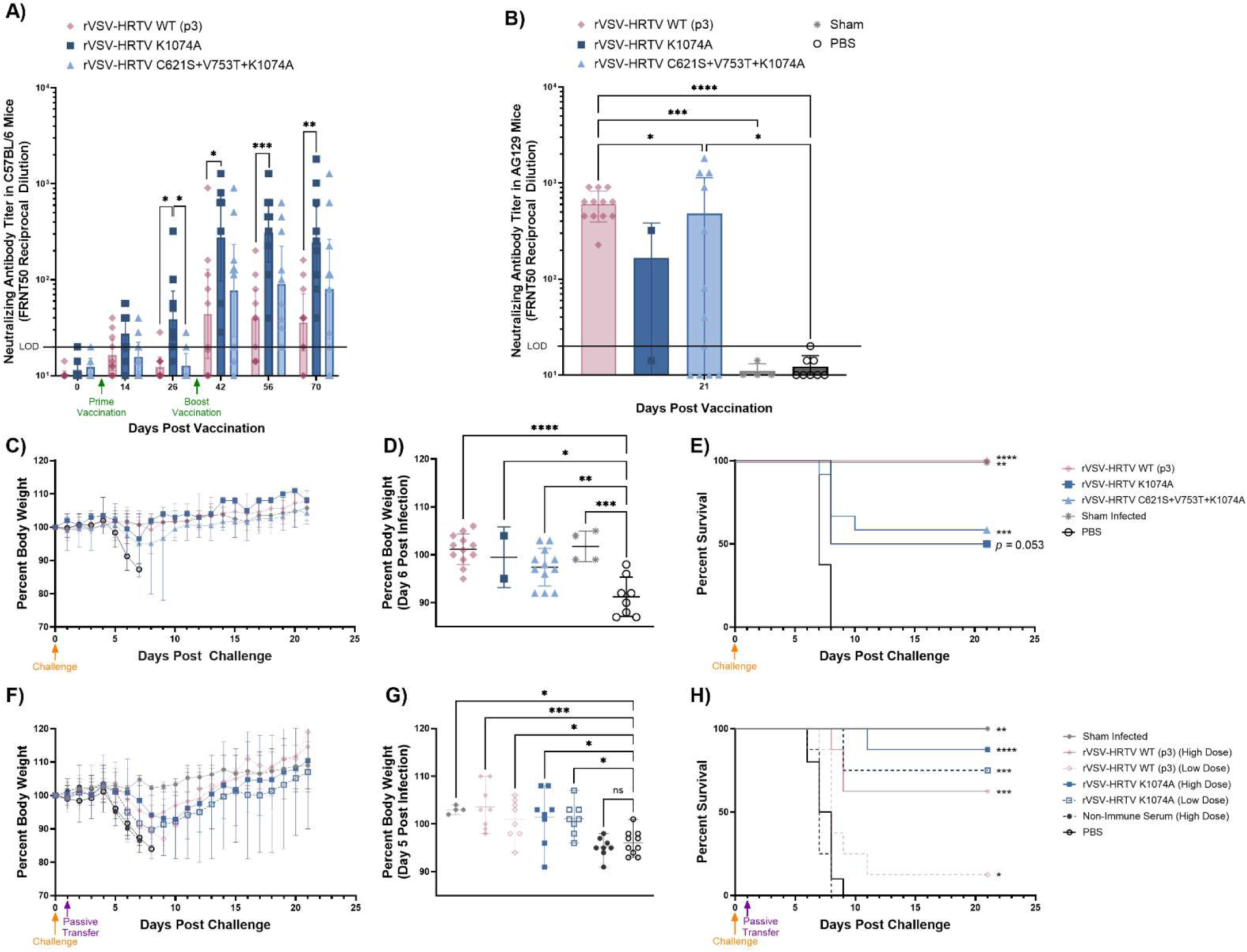
rVSV-HRTV K1074A vaccination results in higher neutralizing antibodies in immunocompetent mice and increased protective efficacy of passive transfer. **(A)** Five male and five female C57BL/6 mice were IP immunized with 10^6^ PFU of rVSV-HRTV WT (p3), K1074A, or C621S+V753T+K1074A at day 0 and day 28. Sera was collected from animals prior to vaccination (day 0), and at 14-, 26-, 42-, 56-, and 70-days post vaccination. Sera was evaluated for neutralizing antibodies against HRTV via FRNT_50_. Horizontal line indicates limit of detection (LOD) of the assay (20) (Two-way ANOVA with Tukey’s multiple comparisons test performed on log normalized data; *, *p* <0.05; **, *p* < 0.01; ***, *p* <0.001). **(B)** Blood was collected from AG129 mice that survived vaccination with 10^2^ PFU of rVSV-HRTV WT (p3), K1074A, or C621S+V753T+K1074A (Figure 6), and neutralizing antibodies against HRTV on day 21 post vaccination were quantified via FRNT_50_. Horizontal line indicates limit of detection (LOD) (20) of the assay (Lognormal ordinary one-way ANOVA with Tukey’s multiple comparisons test; *, *p* <0.05; ***, *p* <0.001; ****, *p* <0.0001). Surviving mice were challenged with MA-HRTV on day 23 post vaccination. **(C)** Weight loss (mean and range) is shown for 21 days post HRTV challenge of immunized mice, and **(D)** compared to the PBS vaccine placebo group at day 6 post challenge (Ordinary one-way ANOVA with Dunnett’s multiple comparisons test; *, *p* <0.05; **, *p* <0.01; ***, *p* <0.001; ****, *p* <0.0001). **(E)** Survival is shown for 21 days post HRTV challenge (Statistical significance of survival was evaluated via Mantel-Cox logrank test compared to mice vaccinated with PBS placebo; *, *p* <0.05; **, *p* <0.01; ***, *p* <0.001; ****, *p* <0.0001). Four male and four female AG129 mice were subcutaneously challenged with MA-HRTV on day 0 and then IP administered serum from mice vaccinated with rVSV-HRTV WT or K1074A one day post challenge. **(F)** Weight loss (mean and range) is shown for 21 days post HRTV challenge, and **(G)** compared to the PBS vaccine placebo group at day 5 post challenge (Ordinary one-way ANOVA with Dunnett’s multiple comparisons test; *, *p* <0.05; ***, *p* <0.001;). **(H)** Survival is shown for 21 days post HRTV challenge (Statistical significance of survival was evaluated via Mantel-Cox logrank test compared to mice vaccinated with PBS placebo; *, *p* <0.05; **, *p* <0.01; ***, *p* <0.001; ****, *p* <0.0001).

To evaluate the protective efficacy of our rVSV-HRTV candidates, we challenged the AG129 mice that survived rVSV-HRTV vaccination (Figure 6B) with 10^2^ CCID_50_ of mouse-adapted HRTV (MA-HRTV) via subcutaneous injection [108]. Blood was collected from the surviving vaccinated mice on day 21 post vaccination so that neutralizing antibody titers could be correlated with survival (Figure 7B). All mice vaccinated with rVSV-HRTV WT (p3) seroconverted, and the average neutralizing titer (640) was very high in these mice. Seroconversion was more variable in the mice vaccinated with rVSV-HRTV K1074A and C621S+V753T+K1074A. Given that the vaccine dose administered was very low (10^2^ PFU), it is possible that these mice did not receive a dose sufficient for immunization or some mice were better able to control early replication of the more attenuated rVSV-HRTV C621S+V753T+K1074A. Nevertheless, the average neutralizing titer of mice vaccinated with rVSV-HRTV K1074A was approximately 167, and the neutralizing titer in the C621S+V753T+K1074A group was 481. Mice were challenged on day 23 post vaccination, and they were evaluated for weight loss (Figure 7C-D), signs of morbidity, and mortality (Figure 7E) for 21 days following challenge. As anticipated, the mice vaccinated with the PBS placebo lost weight rapidly starting at day 4 post challenge and succumbed to viral infection by day 7 post challenge. All the vaccinated groups had elevated survival rates compared to the PBS placebo. All mice vaccinated with rVSV-HRTV WT (p3) survived challenge with minimal weight loss or morbidity. This data correlates with neutralizing antibody titers as all mice both seroconverted and survived challenge. Only two mice survived vaccination with rVSV-HRTV K1074A (Figure 6B), and our data shows that only one of these mice developed neutralizing antibodies (Figure 7B). That mouse survived the MA-HRTV challenge, but the mouse that did not seroconvert did not survive. This trend was similar in the rVSV-HRTV C621S+V753T+K1074A group. Seven of the twelve vaccinated mice developed neutralizing antibodies over the limit of detection, and all but one of these mice survived infection. The mouse that seroconverted but succumbed to infection had a neutralizing antibody titer of 40, so it is possible that this low titer was insufficient for protection.

As seen in Figure 6B, the pathogenicity of rVSV-HRTV K1074A in severely immune-deficient AG129 mice makes it difficult to assess protective efficacy of this vaccine candidate. However, our neutralizing antibody titers from C57BL/6 mice suggest that this vaccine would be the most advantageous in immunocompetent hosts. To directly assess whether the antibody titers elicited by rVSV-HRTV K1074A vaccination in C57BL/6 mice would be protective, we performed a passive transfer study in AG129 mice. We used the serum from the day 70 blood collection to compare the efficacy of passive transfer of pooled serum from rVSV-HRTV WT (p3) (pooled neutralizing titer: 25) or rVSV-HRTV K1074A (pooled neutralizing titer: 253) vaccinated C57BL/6 mice. We chose these samples for evaluation as they had the most disparate phenotypes. Four male and four female AG129 mice per group were subcutaneously challenged with 10^2^ CCID_50_ of MA-HRTV on day 0 and were administered a high or low dose of immune serum via IP injection on day 1 post infection. The high dose consisted of 120 μL undiluted serum, and the low dose consisted of 120 μL (rVSV-HRTV WT) or 110 μL (rVSV-HRTV K1074A) serum diluted by 50%. Additional groups administered 120 μL of non-immune serum or PBS were included as negative controls.

As anticipated, the mice administered non-immune serum or PBS were not protected from infection and therefore lost significant weight and succumbed to HRTV infection between 6 and 9 days post challenge (Figure 7F-H). Administration of serum from rVSV-HRTV WT (p3) vaccinated mice was partially protective. Five of eight mice given the high dose of rVSV-HRTV WT (p3) serum survived infection despite moderate weight loss. The low dose of rVSV-HRTV WT (p3) serum was substantially less effective, and only one mouse in the group survived infection. As we expected based on the neutralizing titers, serum from rVSV-HRTV K1074A was more efficacious than rVSV-HRTV WT (p3). Seven of eight mice that received the high dose of rVSV-HRTV K1074A serum survived infection, and death was significantly delayed in the mouse that succumbed to infection compared to the placebo groups. The low dose of serum from rVSV-HRTV K1074A vaccinated mice was also effective, and six mice from this group survived challenge. These data indicate that antibodies administered via passive transfer are sufficient for protection from lethal HRTV challenge.

## Discussion

SFTSV and HRTV are emerging tick-borne viruses with high case fatality rates, and development of vaccines and therapeutics for these viruses has been prioritized by the WHO and NIAID [28, 29]. Despite this, no vaccines directly targeting HRTV infection have been reported to date, and vaccine candidates for SFTSV have not progressed to more advanced pre-clinical testing. In the case of rVSV-SFTSV, a promising vaccine candidate for SFTSV, one barrier preventing progression to pre-clinical testing is the attenuation of the live virus in cell culture [63, 64, 107]. Previous reports have identified mutations that arise spontaneously in the SFTSV glycoprotein upon passage of rVSV-SFTSV and improve the replication of the virus, but little is understood about the mechanisms of these mutations, their genetic stability, or their effects on antigenicity. These are barriers to clinical implementation of the vaccine and therefore were points of emphasis of our work.

We evaluated the impact of these mutations (C617R, M749T, E982K, and K1071A) on the formation of VSVΔG(SFTSV) pseudotypes, and we discovered that a combination of these substitutions improves the recovery of high titer pseudotypes. More specifically, the K1071A+M749T, K1071A+E982K, and K1071A+C617R+M749T combinations result in nearly a 100-fold increase of pseudotype titers compared to the WT glycoprotein (Figure 1A). This finding alone is important because VSVΔG(SFTSV) pseudotypes have broad utility as diagnostic tools that allow quantification of neutralizing antibodies or identification of potential antiviral drugs in BSL-2 facilities [134]. The pseudotype yield could be further magnified with large scale transfections or concentration of pseudotypes, allowing enhanced downstream applications.

To identify the mechanism(s) responsible for the improved pseudotype titers, we next studied the impact of the substitutions on the biochemical functions of the glycoprotein. Our data suggest that the substitutions impact the stability, fusogenicity, and surface expression of the proteins (Figure 1 B-E). Mutations that increase the fusogenic capabilities or surface expression of glycoproteins have been observed upon passage of rVSVs bearing glycoproteins of other viruses, suggesting that these are relatively common mechanisms of improving recombinant VSV replication that are selected for during passage of diverse rVSV viruses [135, 136]. It is possible that other mechanism(s) share responsibility for the phenotype observed, including altered interactions with host proteins or accumulation of SFTSV glycoproteins in microdomains compatible with VSV proteins, but further work will be necessary to uncover any additional mechanisms at play.

We and others hypothesized the attenuation of rVSV-SFTSV and other rVSVs bearing glycoproteins from the *Bunyaviracetes* class results from mismatches in viral assembly sites [64, 107, 136]. As discussed, SFSTV virions assemble at and acquire their envelopes by budding through the ERGIC and Golgi membranes [118]. This is in contrast to VSV, which assembles at the plasma membrane [83]. Previous studies on SFTSV have demonstrated that Gn and Gc are localized to the Golgi and ER, respectively, when expressed independently, but coexpression of the two glycoprotein components results in transport of Gc from the ER to the Golgi [84]. In this study, we demonstrate that the K1071A substitution in the cytoplasmic tail of SFTSV Gc is sufficient to redirect both Gn and Gc proteins from intracellular compartments to the surface of cells (Figure 2C-D). This finding has been corroborated by additional reports evaluating the cytoplasmic tail of Gc in related bunyaviruses. A 2007 study evaluating the cytoplasmic tail of Uukuniemi virus (UUKV) Gc demonstrated that change of the lysine in the −3 position to alanine (GcK3A) resulted in surface expression of both Gn and Gc [94]. An additional study on Rift Valley Fever virus (RVFV) published in 2014 evaluated the lysine residues found in the Gc cytoplasmic tail and similarly showed that substitution of either lysine residue to alanine resulted in altered localization of Gn and Gc proteins with significant expression at the cell surface [93]. Our study presents data in support of these findings, and we identify the mechanism responsible for the altered localization of SFTSV proteins by showing interaction of the WT SFTSV Gc protein with β’-COP, which is disrupted by the K1071A substitution (Figure 3E). These data support a model in which the COPI retrieval signals in Gc participate in the fine-tuning of Gn/Gc complex distribution, perhaps by targeting the complex for retrograde trafficking between Golgi cisternae or from the Golgi back to the ER. Recruitment of Gc to the Golgi by Gn is thought to require protein-protein interactions between the ectodomains of Gn and Gc. This interaction is thought to bring the cytosolic tails of both proteins into close proximity, allowing the relatively larger Gn cytosolic tail to block binding of the COPI complex to Gc [85, 93]. Intriguingly, the K1071A substitution in the Gc cytosolic tail causes both proteins to leave the Golgi and reach the cell surface. This confirms that COPI is able to bind the cytosolic tail of Gc within the Gn/Gc oligomer to fine-tune the distribution of the complex. It may also indicate that Gc may sterically interfere with the mechanisms responsible for Golgi retention of Gn, though other mechanisms have not been ruled out by our experiments. These results argue for a more complicated explanation for Gn and Gc distribution in cells that is defined by the sum of multiple sorting and trafficking mechanisms. Since the lysine residue at the −3 position of Gc is highly conserved across many bunyaviral families, including *Phenuiviridae, Hantaviridae,* and *Peribunyaviridae* we hypothesize Gc binding to COPI may be more broadly important to the fine-tuning of Gn/Gc protein trafficking of many members of the *Bunyaviricetes* class, and future research will assess this hypothesis directly.

We attempted to launch rVSV-SFTSV viruses with the WT glycoprotein and every combination of mutations described in Figure 1. Despite numerous attempts and extensive troubleshooting, we were unable to launch rVSV-SFTSV with the WT glycoprotein. Given this, we sequenced stocks of our rVSV-SFTSV viruses that were developed for different studies [44], and we found that a mutation encoding the K1071 residue was present in every passage of these stocks dating back to 2013, suggesting that this mutation may have been required for launch or initial spread of our rVSV-SFTSV viruses. In contrast, rVSV-SFSTV encoding the K1071A substitution launched readily and remained genetically stable over subsequent passages. We were able to recover rVSV-SFTSV K1071A, K1071A+C617R, and K1071A+M749T. The only virus we were able to launch without the K1071A substitution was rVSV-SFTSV M749T, as described by Hu et al. Surprisingly, we were unable to launch rVSV-SFTSV C617R+M749T, the substitution combination reported by Hu et al in 2023 [107]. This may be due to differences in the SFTSV strain used, where we employed the HB29 strain of SFTSV but Hu et al. used AH12. Serial passage of our rVSV-SFTSV viruses that were successfully rescued revealed that only rVSV-SFTSV K1071A was genetically stable. All other viruses accumulated additional mutations as they were passaged (Figure 4A). We evaluated the replication kinetics of all the rVSV-SFTSV mutant viruses we were able to launch. As anticipated, rVSV-SFTSV M749T did not replicate very efficiently, but the addition of the K1071A substitution improved replication significantly. However, we were surprised to find that rVSV-SFTSV K1071A replicated very efficiently on its own, and this replication was not markedly improved by the addition of other mutations (Figure 4B). Similarly, we were surprised to see that glycoproteins with only the K1071A substitution were incorporated onto VSV virions much more efficiently than K1071A+C617R or K1071A+M749T (Figure 4E). Given that there is no significant difference in surface expression between these three viruses, the difference in incorporation is striking. This may also be a result of increased fusogenicity as the glycoproteins may be triggered to adopt the post fusion conformation more readily, hindering incorporation into virions.

Comparison of SFTSV and HRTV glycoproteins revealed cognate residues in HRTV. To evaluate the effects of mutation of these residues in HRTV, we launched rVSV-HRTV viruses with WT glycoproteins and glycoproteins with the K1074A or C621S+V753T+K1074A substitutions. Overall, our rVSV-HRTV data were similar to rVSV-SFTSV. We found that only rVSV-HRTV K1074A was genetically stable, and rVSV-HRTV WT and rVSV-HRTV C621S+V753T+K1074A acquired additional mutations during serial passage (Figure 5A). We also found that rVSV-HRTV K1074A replicated efficiently, and replication was not improved by adding the C621S+V753T substitutions (Figure 5B). We confirmed that glycoproteins in rVSV-HRTV K1074A infected cells were expressed on the cell surface at significantly higher levels than in rVSV-HRTV WT or rVSV-HRTV C621S+V753T+K1074A infected cells (Figure 5D). Furthermore, glycoproteins from rVSV-HRTV K1074A and rVSV-HRTV C621S+V753T+K1074A were incorporated into VSV virions at much higher levels than the WT glycoprotein (Figure 5E).

Taken together, these data indicate that the K1071A and K1074A substitutions that alter a non-canonical COPI binding site in SFTSV and HRTV Gc, respectively, are sufficient to improve performance of rVSV *in vitro*. The addition of C617R/C621S or M749T/V753T substitutions in the launched viruses decreases the genetic stability of the rVSV, allowing the accumulation of additional mutations during serial passage. The effect of these mutations that arise spontaneously are unknown and they may impair the immunogenicity of the glycoprotein payload or have other unexpected deleterious consequences. Overall, the genetic instability would inhibit progression of the vaccine into the clinic. It is therefore favorable to produce viruses with a targeted K1071A/K1074A genetic alteration as they are genetically stable and have a known biological outcome. Additionally, the K1071A/K1074A substitutions lie in the cytoplasmic tail of the glycoprotein, and there is therefore much less concern that they impact an important antigenic site or interfere with B cell epitopes. Finally, replication of rVSV-SFTSV K1071A and rVSV-HRTV K1074A is efficient and not readily improved by other mutations. Our data suggest that the other substitutions, particularly M749T, increase the fusogenicity of the glycoproteins, resulting in an augmented capacity for cell-cell spread (Figure 1D-E, 4C, and 5C). However, this would negatively affect effective vaccine development as it is well understood that the prefusion conformation of viral glycoproteins are preferred for induction of neutralizing antibodies, but the conformational changes that occur during fusion alter the antigenicity of the protein [137–139].

Our data also support the conclusion that rVSV-HRTV K1074A outperforms the other viruses *in vivo,* resulting in an improved vaccine candidate. We found that rVSV-HRTV K1074A replicates significantly more efficiently in mice, as evidenced by the rapid weight loss and mortality in immunodeficient AG129 mice injected with rVSV-HRTV K1074A. In contrast, AG129 mice that are dosed with rVSV-HRTV WT (p3) and rVSV-HRTV C621S+V753T+K1074A lose only a moderate or mild amount of weight and recover quickly from infection (Figure 6A-B). Supporting increased *in vivo* replication of the K1074A virus, VSV-N RNA is detected in sera of rVSV-HRTV K1074A infected mice while VSV-N RNA is below the limit of detection of our assay in sera from mice infected with rVSV-HRTV WT (p3) and rVSV-HRTV C621S+V753T+K1074A. This improvement in replication is also suggested by the immunogenicity data from immunocompetent C57BL/6 mice, where we show that vaccination with rVSV-HRTV K1074A results in induction of higher neutralizing antibody titers than vaccination with either of the other two candidates (Figure 7A). C57BL/6 mice were evaluated for signs of morbidity, including hunched posture, unkempt fur, and decreased activity. We did not observe any signs of clinical illness resulting from rVSV-HRTV K1074A, or any of the other rVSV-HRTV variants, in C57BL/6 mice, which indicates that there is not an increased risk of this live vaccine in immune competent hosts. We found that vaccination with rVSV-HRTV C621S+V753T+K1074A results in lower neutralizing antibody titers at every time point, although this difference did not meet statistical significance except for on day 26 post vaccination. This difference in neutralizing antibody titers may result from impaired *in vivo* replication, but it is also possible that the mutations that appear to increase the fusogenicity of the glycoprotein allow the protein to undergo conformational changes more readily, resulting in a post fusion structure that is less immunogenic, as described above.

We evaluated protective efficacy of our rVSV-HRTV vaccine candidates in the severely immunocompromised AG129 mouse model. Immunocompromised mice deficient in interferon responses are not ideal for evaluation of vaccine responses due to the critical role of interferon in the development of antibody and T cell responses [140, 141]. However, there are no acceptable immunocompetent models for HRTV infection, which limits our ability to assess vaccine efficacy. Immunocompetent C57BL/6 mice do not develop viremia or any other symptoms of HRTV viral infection [142]. *Ifnar^-/-^*mice are deficient only in interferon α and β receptors and are therefore slightly less immunocompromised, but they are only susceptible to HRTV infection at exceptionally high doses delivered intraperitoneally, which is prohibitive for many studies [143].

Despite the severe pathogenicity of rVSV-HRTV K1074A, our challenge study in AG129 mice provided critical data (Figure 7B-E). We observed universal seroconversion with high neutralizing antibody titers and complete protection against HRTV after vaccination with rVSV-HRTV WT (p3). Responses to rVSV-HRTV C621S+V753T+K1074A were more variable; seven of the twelve vaccinated mice developed detectable neutralizing antibodies with a wide range of titers (FRNT_50_ reciprocal dilution of 40 to 1810). Of these seven mice, all but the mouse with the lowest neutralizing titer (40) survived HRTV challenge. These data suggest that neutralizing antibodies may be an important correlate of protection for HRTV infection, as is the case for SFTSV [48–53]. However, one mouse vaccinated with rVSV-HRTV C621S+V753T+K1074A did not develop detectable neutralizing antibodies but survived lethal challenge, implying that cellular immunity or non-neutralizing antibodies can play a role in protection from HRTV infection. Supporting this idea, protection in the absence of high titer neutralizing antibodies has been previously described in a study evaluating the efficacy of rVSV-SFTSV in cross-protection against HRTV [64].

To directly evaluate the role of antibodies in protection from HRTV challenge, as well as to more effectively demonstrate the improved efficacy of our rVSV-HRTV K1074A vaccine candidate, we performed a passive transfer study. In this experiment, we pooled serum from C57BL/6 mice vaccinated with rVSV-HRTV WT (p3) (neutralizing titer: 25) or rVSV-HRTV K1074A (neutralizing titer: 253) and administered high or low doses of serum to AG129 mice that had been challenged with MA-HRTV (Figure 7F-H). We found that serum from rVSV-HRTV WT (p3)-vaccinated mice was moderately effective at the high dose (62.5% survival), but efficacy was dramatically decreased at the low dose (12.5% survival). Serum from mice vaccinated with rVSV-HRTV K1074A was efficacious at both a high and low dose, with 87.5% survival and 75% survival, respectively. There are several important conclusions from these data. First, this indicates that vaccination with rVSV-HRTV K1074A in immunocompetent mice produces more potent antibody responses with higher protective efficacy than rVSV-HRTV WT (p3). Second, this provides additional support for the role of neutralizing and non-neutralizing antibodies in protection from HRTV infection. We see a clear dose-dependent and neutralizing titer-dependent response in mice receiving passive transfer. However, some protection was still exhibited in mice administered rVSV-HRTV WT (p3)-vaccinated serum. Given the low initial neutralizing titer of this serum and the dilution within the recipient mouse’s circulatory system, it is unlikely that there would be an appreciable neutralizing titer in the recipient mice. This indicates that non-neutralizing antibodies may play a role in protection. Finally, we show that transfer of immune serum or development of monoclonal antibodies may be a possible therapeutic consideration in infected patients, but more work is required to understand the potential efficacy of this treatment.

Overall, we identify a substitution (K1071A) in the cytoplasmic tail of the SFTSV Gc glycoprotein that is sufficient to redirect viral proteins from intracellular compartments to the cell surface, where they can be more readily incorporated onto budding VSV particles. This phenomenon occurs via alteration of a noncanonical COPI binding signal in the cytoplasmic tail of the protein. To our knowledge, this is the first study directly assessing the role of COPI binding in cellular localization of a bunyavirus. More work needs to be done to generalize this finding to other viruses in the *Bunyaviracetes* class, but conserved lysine residues have been identified in the cytoplasmic tails of *Phenuiviridae, Hantaviridae,* and *Peribunyaviridae* families. A similar substitution of lysine to alanine in the cytoplasmic tails of bunyaviruses from these viruses may allow enhanced incorporation of glycoproteins onto VSV, both for production of pseudotypes and for generation of rVSV, and this possibility will be explored in future studies. More broadly, our studies should inform other vaccine platforms, such as mRNA-LNP, where increased surface expression of bunyaviral glycoproteins would likely increase vaccine efficacy. We show that the K1071A substitution improves replication of rVSV-SFSTV *in vitro,* and a K1074A substitution in the cognate site of rVSV-HRTV has similar effects. *In vivo* studies involving vaccination of mice with rVSV-HRTV K1074A demonstrate improved replication in animal systems, which results in increased immunogenicity in immunocompetent C57BL/6 mice. To date, this is the first report of a vaccine directly targeting HRTV infection and of the use of HRTV immune sera to provide protection. Our data support the use of rVSV-HRTV K1074A as a potential vaccine candidate as it induces high titers of neutralizing antibodies that protect severely immunocompromised mice from lethal challenge when administered via passive transfer. Future studies should explore whether rVSV-HRTV vaccination or immune sera can offer cross-protection from SFTSV.

## Acknowledgements

We are grateful for the technical support provided by Dionna Scharton. This work was supported by the Martin and Pamela Winter Infectious Disease Fellowship from the Institute for Infectious and Zoonotic Diseases of the University of Pennsylvania School of Veterinary Medicine and the National Institute of Allergy and Infectious Diseases, National Institutes of Health (R21AI142638, R01AI152236, T32AI055400, T32AI070077, R41AI174426, and HHSN272201700041I).

